# Decreasing peptide deformylase activity is a beneficial strategy for increasing formaldehyde resistance in *Methylobacterium extorquens*

**DOI:** 10.64898/2026.04.16.718930

**Authors:** Chandler N. Hellenbrand, Tyler Miller, Elias I. Kemna, Eric L. Bruger, Zachary T. Hying, Jannell V. Bazurto

## Abstract

Formaldehyde is a highly toxic metabolite that can cause extensive damage to DNA and proteins, and strategies to mitigate formaldehyde toxicity are poorly understood. Methylotrophic bacteria, such as *Methylobacterium extorquens*, thrive on one-carbon compounds as sole sources of carbon and energy. These organisms are excellent models for discovering formaldehyde stress response systems because formaldehyde is an obligate intermediate in their central carbon metabolism. Here, we characterize an evolved *def* allele (*def^evo^*) that increases formaldehyde resistance in *M. extorquens*. The *def* gene encodes peptide deformylase (PDF, EC:3.5.1.88), an enzyme that contributes to protein processing by removing the formyl group from *N-*formylmethionine (fMet) on nascent peptides. The *def^evo^*allele has a single missense mutation that decreases PDF activity both *in vitro* and *in vivo*. Transcriptomic analysis of the *def^evo^* strain indicates there are pleiotropic effects of this mutation and a differential response to formaldehyde stress. We investigate possible mechanisms for the *def^evo^* mutant’s increased resistance to formaldehyde, including mitigation of formaldehyde-induced protein stress and altered membrane physiology. We find that the *def^evo^* allele selectively alleviates exogenous, but not endogenous, formaldehyde stress and identify a tradeoff in heat shock resistance. This study reports the first observation of lowered PDF activity benefiting a cellular physiological phenotype. Our work indicates that altered protein metabolism can mitigate the toxic effects of formaldehyde and furthers our understanding of the strategies that can protect cells from formaldehyde-induced damage.

**Importance:** Formaldehyde is a toxic chemical that can damage essential molecules inside of cells, yet all organisms inevitably produce it during normal metabolism. Despite its ubiquity, our understanding of strategies for how cells navigate formaldehyde toxicity is incomplete. This study focuses on *Methylobacterium extorquens*, which naturally generates high levels of formaldehyde as part of its growth on simple carbon compounds. We show herein that a single genetic change, which slows down how newly made proteins are processed during translation, can unexpectedly improve the bacterium’s ability to resist formaldehyde stress. Further, we show that this single change has numerous effects on the cell, many of which may contribute to formaldehyde resistance.

## Introduction

Formaldehyde is an unavoidable byproduct of metabolic reactions and cellular processes in all organisms. At elevated concentrations, this simple aldehyde can damage DNA, proteins, lipid membranes, and other biologically important molecules (1–4). The universal toxicity of formaldehyde is attributed to its electrophilic nature, making it highly reactive with nucleophilic structures, such as the amine and thiol groups that are prevalent on nucleotides and amino acids (5). This renders DNA and protein molecules highly susceptible to formaldehyde-induced modifications that can damage these critical cellular components (6, 7).

Formaldehyde’s reactivity with amino acids makes it an effective proteotoxic stressor (3). Amines and thiols react with formaldehyde to form unstable *N*-hydroxymethyl and *S*-hydroxymethyl adducts, respectively (Fig. 1A, B). While this initial reaction is reversible, the products can undergo secondary reactions that result in permanent modifications (4, 8–10). *N*- and *S-*hydroxymethyl adducts can react with other primary amines to form covalently crosslinked methylene bridges. For instance, cysteine residues can cyclize through an intramolecular methylene bridge, forming a thiazolidine ring structure (also called thioproline)(11).

**Figure 1:**
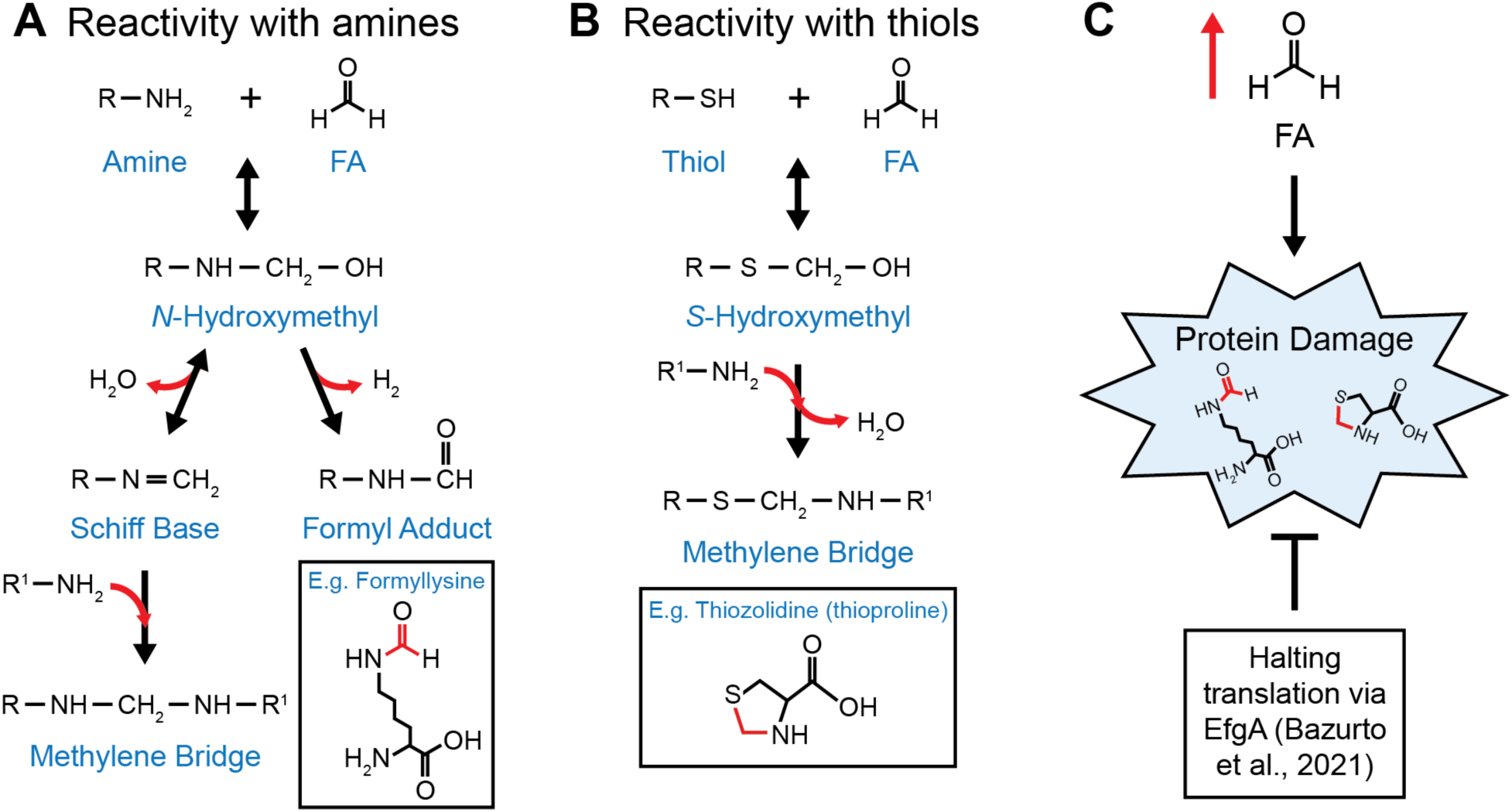
Formaldehyde is an effective proteotoxic stressor. Formaldehyde (FA) reactivity with A) amines and B) thiols. When proteins are exposed to formaldehyde, amino acids with these nucleophilic groups (e.g., lysine, cysteine) react with formaldehyde and cause protein damage (e.g. formyllysine, thioproline). C) *M. extorquens* can respond to elevated formaldehyde and prevent protein damage by halting translation through EfgA (19).

Due to the ubiquitous nature of formaldehyde, nearly all organisms have evolved formaldehyde detoxification pathways (4, 12–14). Many of these detoxification pathways involve the conversion of formaldehyde to a nontoxic metabolite, such as formate, that can then be assimilated into other metabolic pathways. However, beyond detoxification, alternative mechanisms used to sense, respond to, and mitigate formaldehyde stress are less understood, including processes such as removal, sequestration, physiological reconfiguration, and repair (15). Methylotrophs, such as the Gram-negative bacterium *Methylobacterium extorquens*, present unique opportunities to study formaldehyde stress response mechanisms. *M. extorquens* is a facultative methylotroph and can use single-carbon (C1) compounds like methanol as a sole source of carbon and energy. The first step to utilizing methanol is the oxidation to formaldehyde by periplasmic methanol dehydrogenase. Once in the cytosol, formaldehyde is catalytically condensed with the C1 carrier, dephospho-tetrahydromethanopterin (dH4MPT) by the formaldehyde activating enzyme (Fae), acting as both a detoxification and an oxidation pathway (16). Due to its unique metabolism, *M. extorquens* has a high tolerance to formaldehyde and can endure intracellular concentrations of up to 2 mM before losing viability (17–20).

While methylotrophic pathways for the consumption and assimilation of formaldehyde have been known for decades, the mechanisms by which *M. extorquens* and other methylotrophs sense and respond to formaldehyde stress are just beginning to be uncovered. Although formaldehyde is an obligate intermediate during methylotrophy, wild-type *M. extorquens* is unable to grow on formaldehyde as a sole carbon and energy source. However, certain evolved mutations have been shown to increase formaldehyde resistance enough for *M. extorquens* to grow directly on formaldehyde. Prominent among these are loss-of-function mutations in *efgA.* EfgA is a recently identified formaldehyde sensor that appears to have a role in protecting cells from formaldehyde-induced damage (19–21). When *in vivo* formaldehyde levels are elevated, EfgA binds free formaldehyde and arrests growth and translation. The abrupt cessation of protein synthesis acts to prevent formaldehyde from damaging susceptible molecular targets before the cell can detoxify the highly reactive metabolite (Fig. 1C). Intriguingly, while cells lacking *efgA* are more resistant to treatment with exogenous formaldehyde, this resistance comes at the cost of a defective transition to methylotrophic growth, where they fail to maintain homeostasis of endogenously generated formaldehyde and consequently lose viability (20).

Additional evolved mutations in the *def* gene (*Mext_1636*) suggest that modulating protein metabolism is a key factor that allows *M. extorquens* to tolerate high concentrations of formaldehyde. *def* encodes peptide deformylase (PDF), an essential ribosome-associated enzyme involved in protein maturation. PDF removes the formyl group from the first amino acid, *N-*formylmethionine (fMet), on nascent peptides during bacterial translation (22–24). While the catalytic activity of PDF and its role in protein maturation are well-characterized, the physiological significance of PDF and its fMet-containing substrates remains poorly understood. It is postulated that fMet has a role in protein quality control (25, 26), and PDF activity impacts various aspects of bacterial physiology. In *E. coli*, PDF inhibition increases the translation of proteins related to transcription and translation, compromises membrane integrity, and disrupts redox metabolism (27).

An evolved *def* allele (*def^evo^*), which encodes a single missense mutation in PDF (V54G), allows *M. extorquens* to grow on formaldehyde as a sole carbon source. Here, we show that this mutation causes a ∼60% reduction in PDF activity and increases the abundance of fMet present on newly synthesized proteins. The *def^evo^* allele confers general formaldehyde resistance, but it does not do so through the mitigation of proteotoxicity or by decreasing formaldehyde uptake. Transcriptomic analysis of the *def^evo^* mutant revealed pleiotropic effects on the transcriptome, including a differential response to formaldehyde treatment. These findings provide the first example of decreased PDF activity leading to improved growth of a bacterial strain and raises questions on how protein maturation affects the ability to tolerate a universally encountered and toxic metabolite.

## Results

### Evolved *def* alleles have increased formaldehyde resistance

Our previous work demonstrated that single evolved alleles in *def*, which encodes peptide deformylase (PDF), the ribosome-associated enzyme responsible for removing the formyl group from *N-*formylmethionine (fMet) on nascent bacterial peptides, allow wild-type (WT) *M. extorquens* to grow on formaldehyde as a sole carbon and energy source (19). A mutant harboring one of these evolved *def* alleles encoding a V54G substitution, *def^evo^*, was grown on succinate with varying concentrations of formaldehyde to test if this mutant’s ability to grow on formaldehyde translated to an increase in formaldehyde resistance (Fig. 2A). WT and *def^evo^* strains grow comparably in media containing 3.5 mM succinate and no supplemented formaldehyde, but the *def^evo^* mutant has a shorter lag phase than WT as the formaldehyde concentration in the media increases. For instance, the *def^evo^* mutant begins exponential growth after just 20 hr when treated with 4 mM formaldehyde, while WT remains in lag phase for 56 hr (Fig. 2A). This decreased sensitivity to formaldehyde is consistent with other reported evolved alleles that allow *M. extorquens* to grow on formaldehyde, namely loss-of-function mutations in *efgA* (Fig. S1).

**Figure 2:**
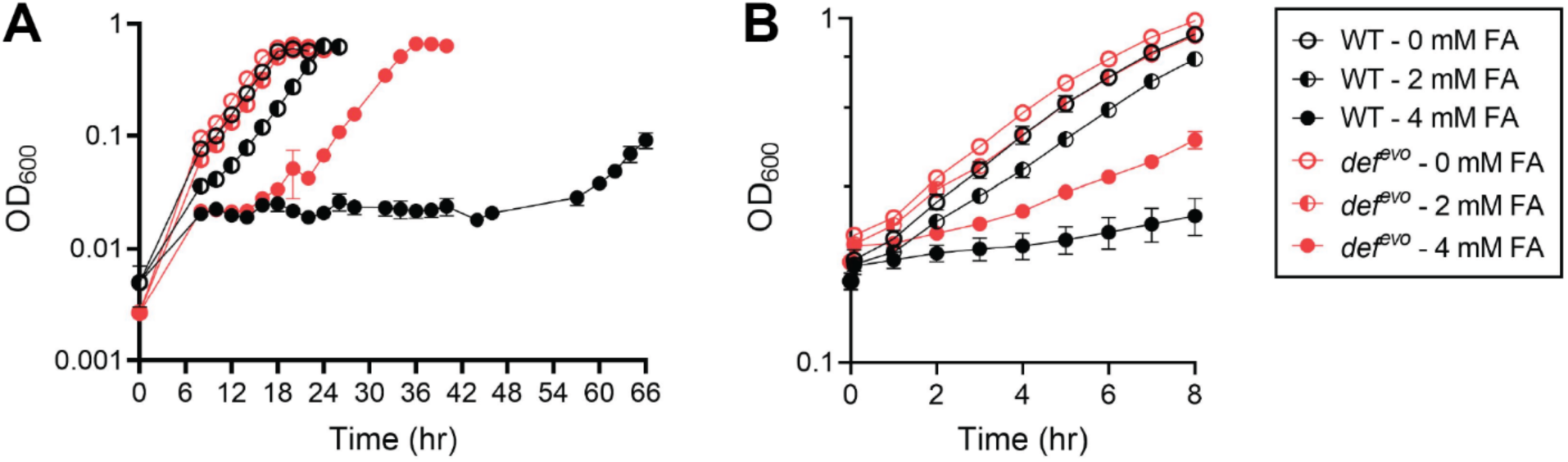
The *def^evo^* mutant has increased formaldehyde resistance compared to WT. A) WT and *def^evo^* strains were grown in MP with 3.5 mM succinate and 0, 2, or 4 mM formaldehyde. B) WT and *def^evo^* were grown in MP with 15 mM succinate and exposed to 0, 2, or 4 mM formaldehyde in early exponential phase (OD600=0.2). WT shows a longer lag in growth than *def^evo^* under all conditions with formaldehyde stress. The error bars represent the standard deviation of the mean for three biological replicates.

A similar pattern is seen when WT and *def^evo^* strains are exposed to formaldehyde in early exponential phase (Fig. 2B). Strains were grown to an OD600 of 0.2 in 15 mM succinate and then shocked with either 2 mM or 4 mM formaldehyde. The *def^evo^* strain was less inhibited than WT at both concentrations. We conclude that the single V54G substitution in PDF confers increased formaldehyde resistance in *M. extorquens*, and this phenotype likely contributes to the *def^evo^* strain’s ability to grow on formaldehyde.

### The evolved *def^evo^* allele has decreased peptide deformylase activity *in vivo*

We next investigated how the V54G substitution impacts PDF activity. The *def^evo^* strain can grow on formaldehyde as a sole carbon source while WT cannot (19). When the wild-type *def* gene is ectopically expressed in the *def^evo^* mutant, the wild-type phenotype is restored, and the complemented strain can no longer grow on formaldehyde (Fig. 3A). The recessive nature of the evolved allele suggests that the PDF^V54G^ variant has reduced deformylase activity *in vivo*, as wild-type PDF compensates for its diminished function. When grown on succinate alone (Fig. 3A), WT and *def^evo^* grow indistinguishably from one another with nearly identical growth rates (Table 1). An actinonin minimum inhibitory concentration (MIC) assay further supported that the PDF^V54G^ variant has less *in vivo* activity (Fig. 3B). Actinonin is an antibiotic that directly inhibits PDF by mimicking its substrate, N-formylmethionine, creating competitive inhibition and occluding the essential function of PDF (28). WT and *def^evo^* strains were inoculated into media containing 0-100 µg/mL actinonin to test their sensitivity to the antibiotic. We found that the *def^evo^* mutant was more sensitive to actinonin at every concentration tested (Fig. 3B). Less actinonin is needed to substantially impair the growth of the *def^evo^* strain compared to WT, further indicating that the evolved PDF^V54G^ variant has less *in vivo* deformylase activity.

**Figure 3:**
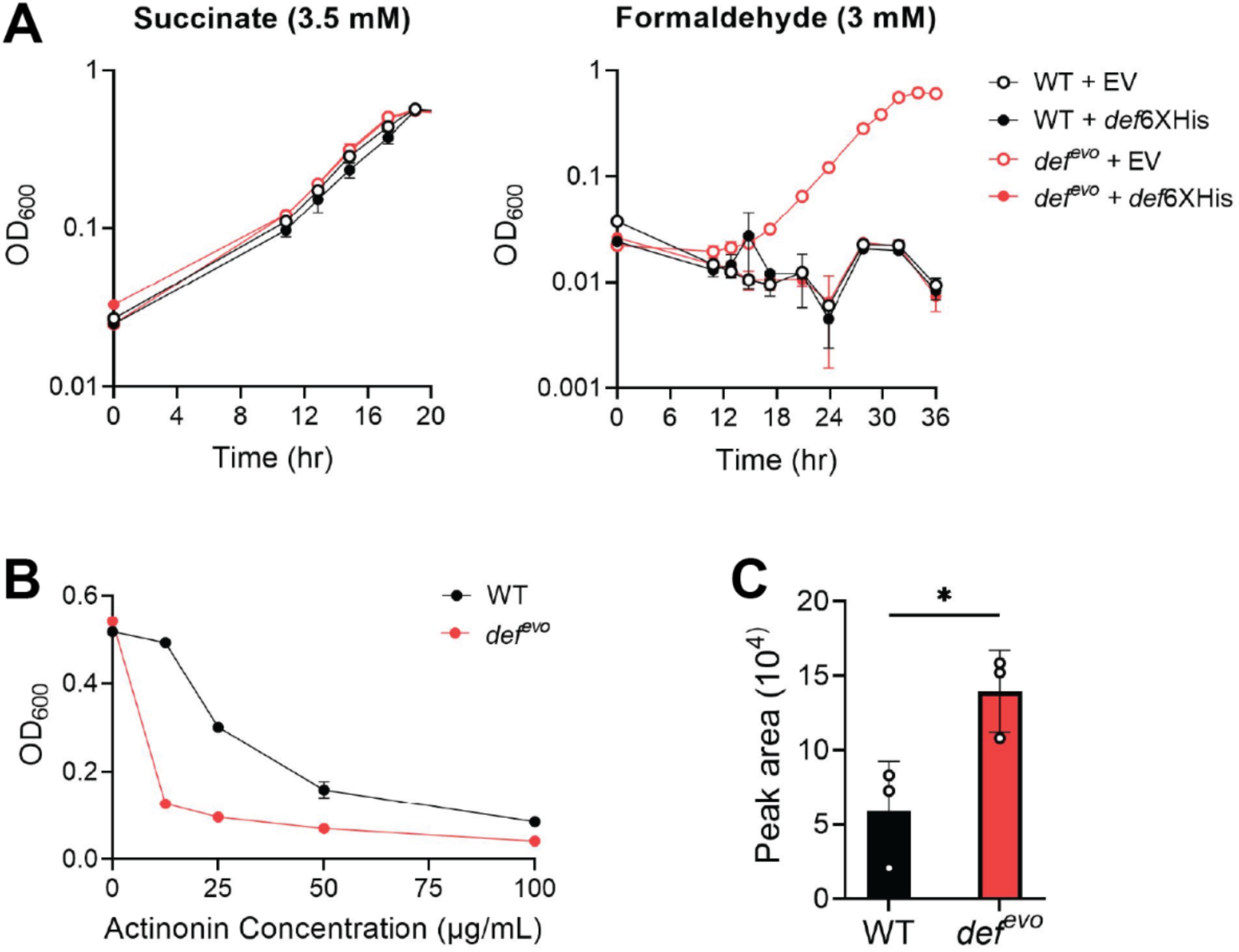
The *def^evo^* strain has less *in vivo* peptide deformylase activity than WT. A) WT and *def^evo^* strains with *def^6XHis^* (pCH14) or an empty pCH07 vector were grown on MP media with 3.5 mM succinate (left) or 3 mM formaldehyde (right). In succinate, all strains grow comparably with similar growth rates. When formaldehyde is the sole carbon source, expressing *def^6XHis^* in the *def^evo^* mutant restores the wild-type phenotype of no growth. B) The *def^evo^* mutant is more sensitive to PDF-inhibitor actinonin. Strains were grown on MP with 15 mM methanol and 0-100 µg/mL actinonin. Optical density was recorded after 30 hr of growth. C) The *def^evo^*mutant has increased levels of fMet, as measured by mass spectrometry. \**P* < 0.05. All error bars represent the standard deviation of the mean for three biological replicates.

**Table 1.**
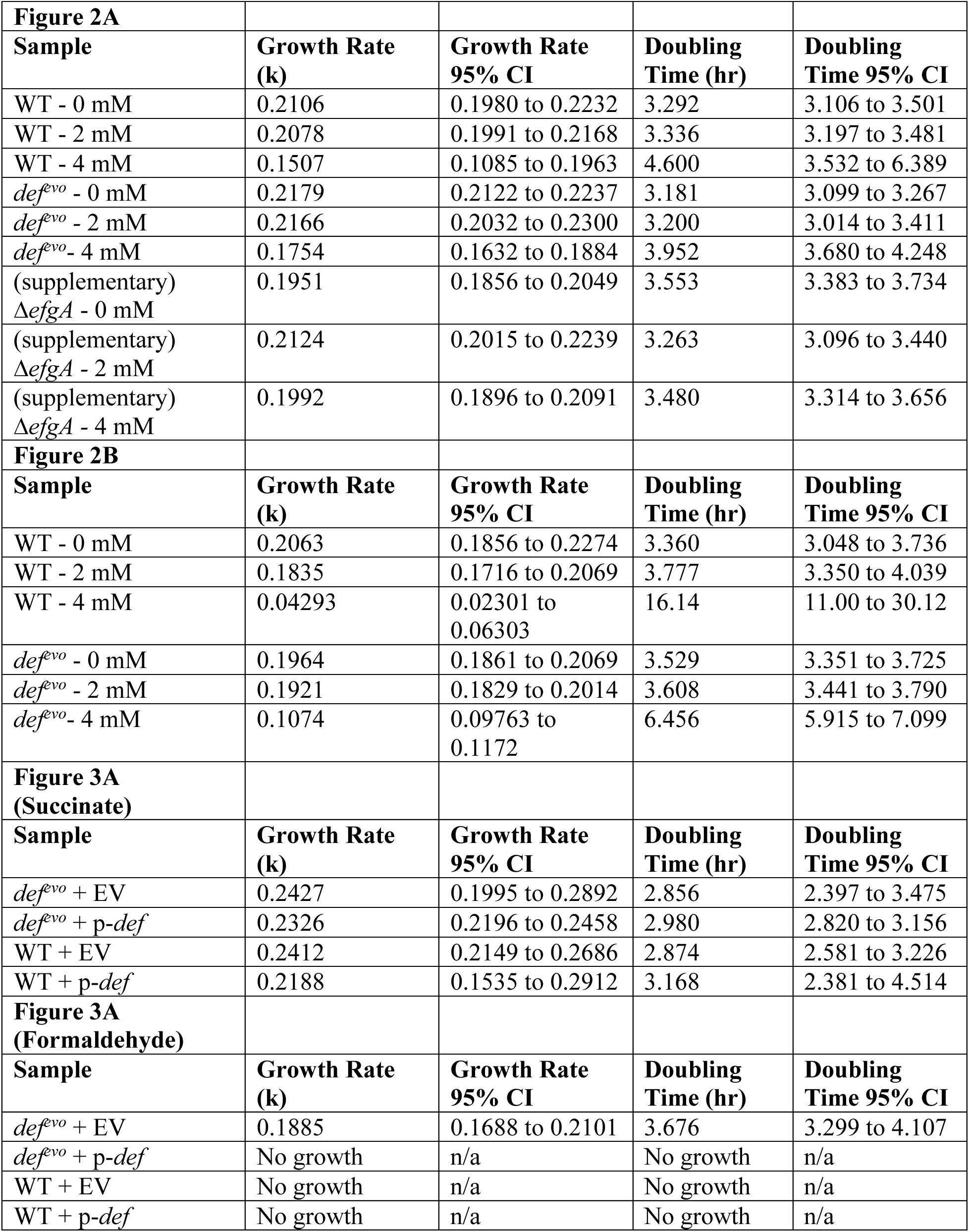
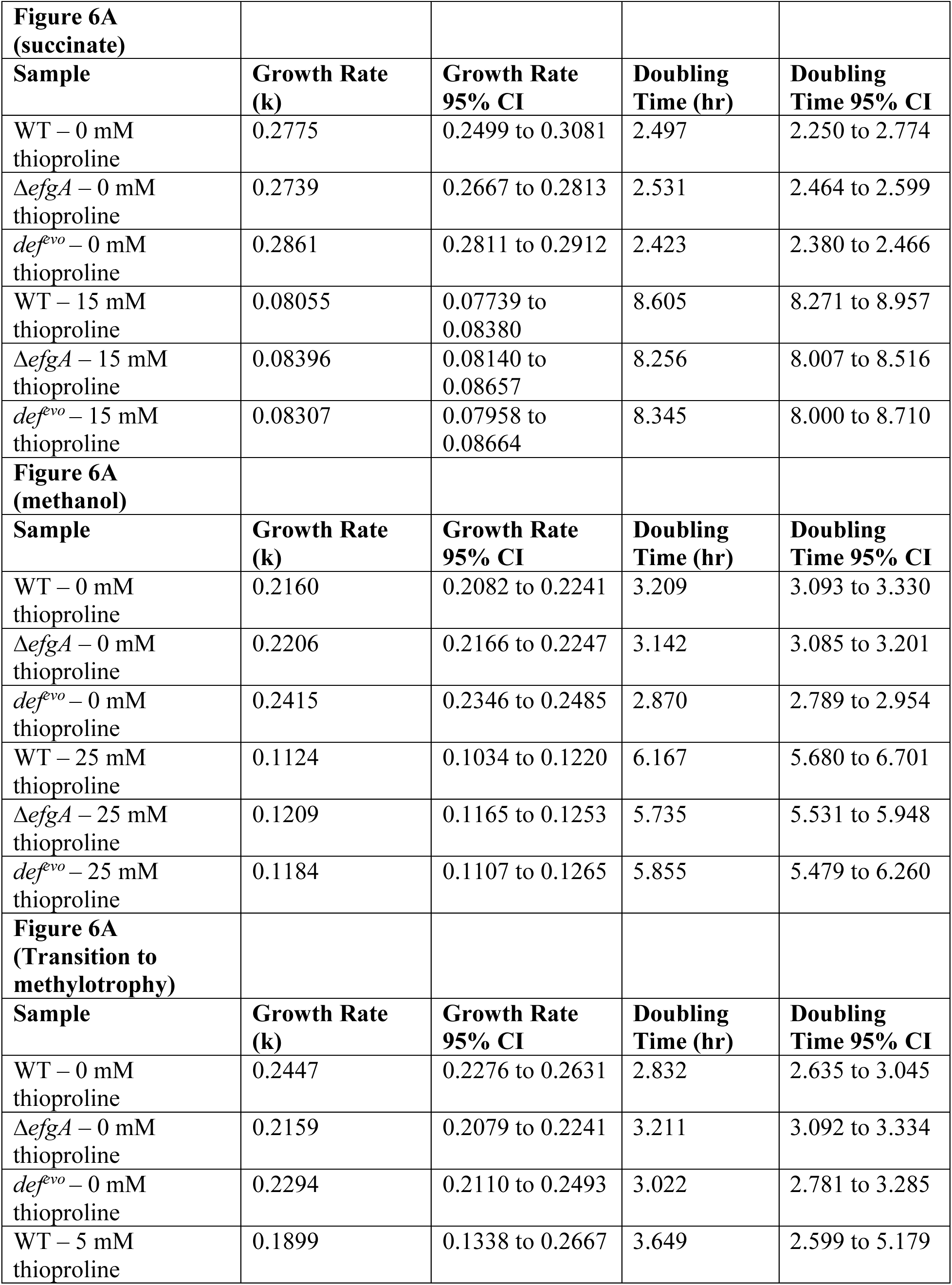
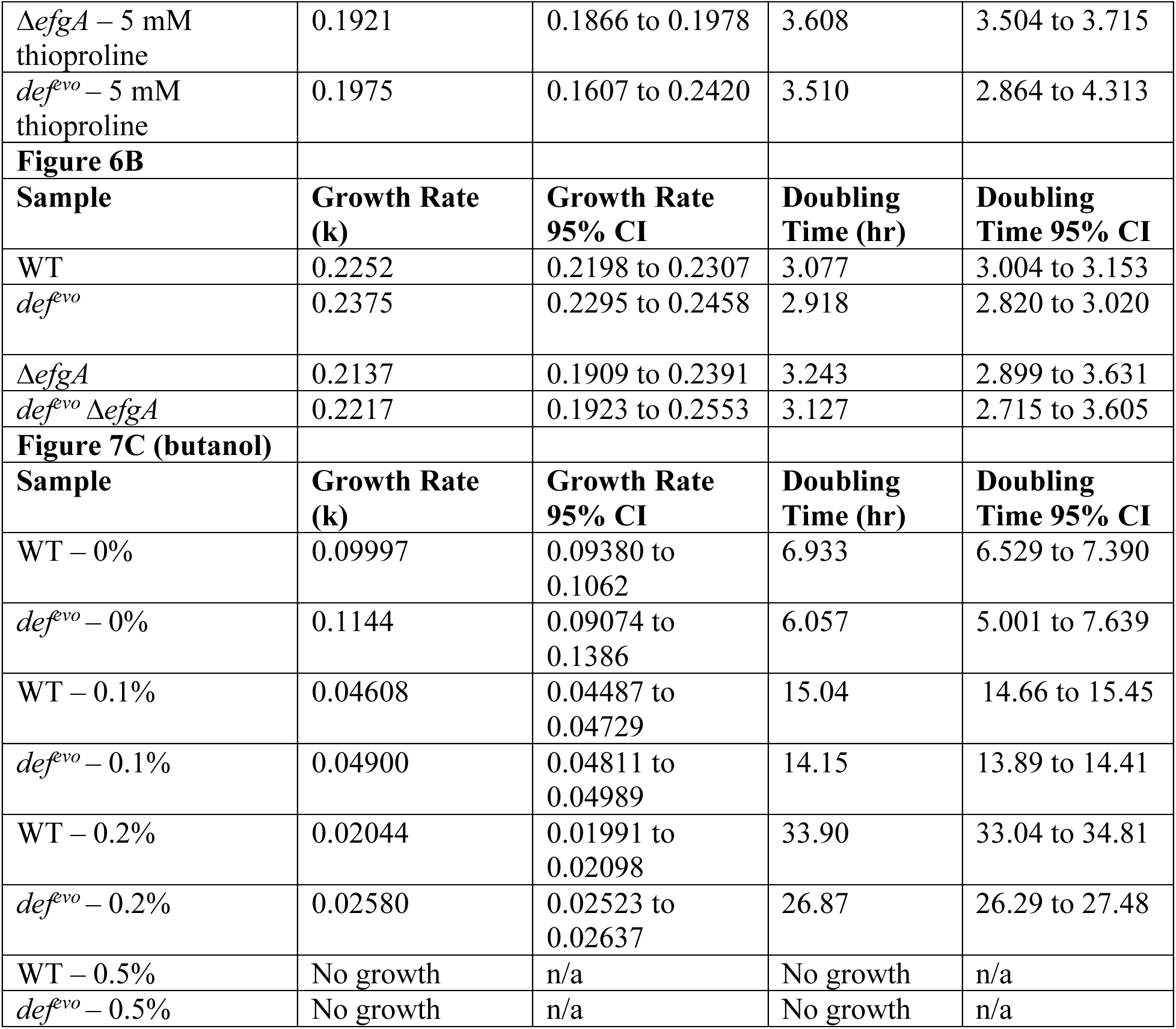
Growth rates.

### The PDF^V54G^ variant enzyme has reduced catalytic activity *in vitro*

We expressed and purified recombinant PDF^WT^ and PDF^V54G^ in *E. coli* to investigate their relative *in vitro* activity. PDF activity can be measured with a colorimetric assay using the peptide substrate N-formyl-methionylleucyl-p-nitroanilide (fML-pNA) coupled with *Aeromonas* aminopeptidase (AAP)(29–31). When incubated together, PDF deformylates fML-pNA, exposing the N-terminus of the substrate. The substrate is then recognized by AAP, which hydrolyzes the substrate into its components and releases the chromogenic *p-*nitroaniline (ε405 nm = 10,600 M^-1^ cm^-1^). This assay was performed with purified PDF^WT^ and PDF^V54G^ with substrate concentrations ranging from 0-400 mM (Fig. 4A). PDF^WT^ was found to have a KM of 80.81 ± 4.44 µM and a kcat of 12.74 ± 0.34 s^-1^. The kinetic values for PDF activity found here are comparable to those found in other bacteria (29, 31, 32). The PDF^V54G^ variant had a similar KM of 78.76 ± 3.05 µM, but its kcat was decreased to 5.39 ± 0.13 s^-1^. This indicates that the PDF^V54G^ enzyme has the same affinity for the fML-pNA substrate as PDF^WT^ but has a 58% decrease in turnover number.

**Figure 4:**
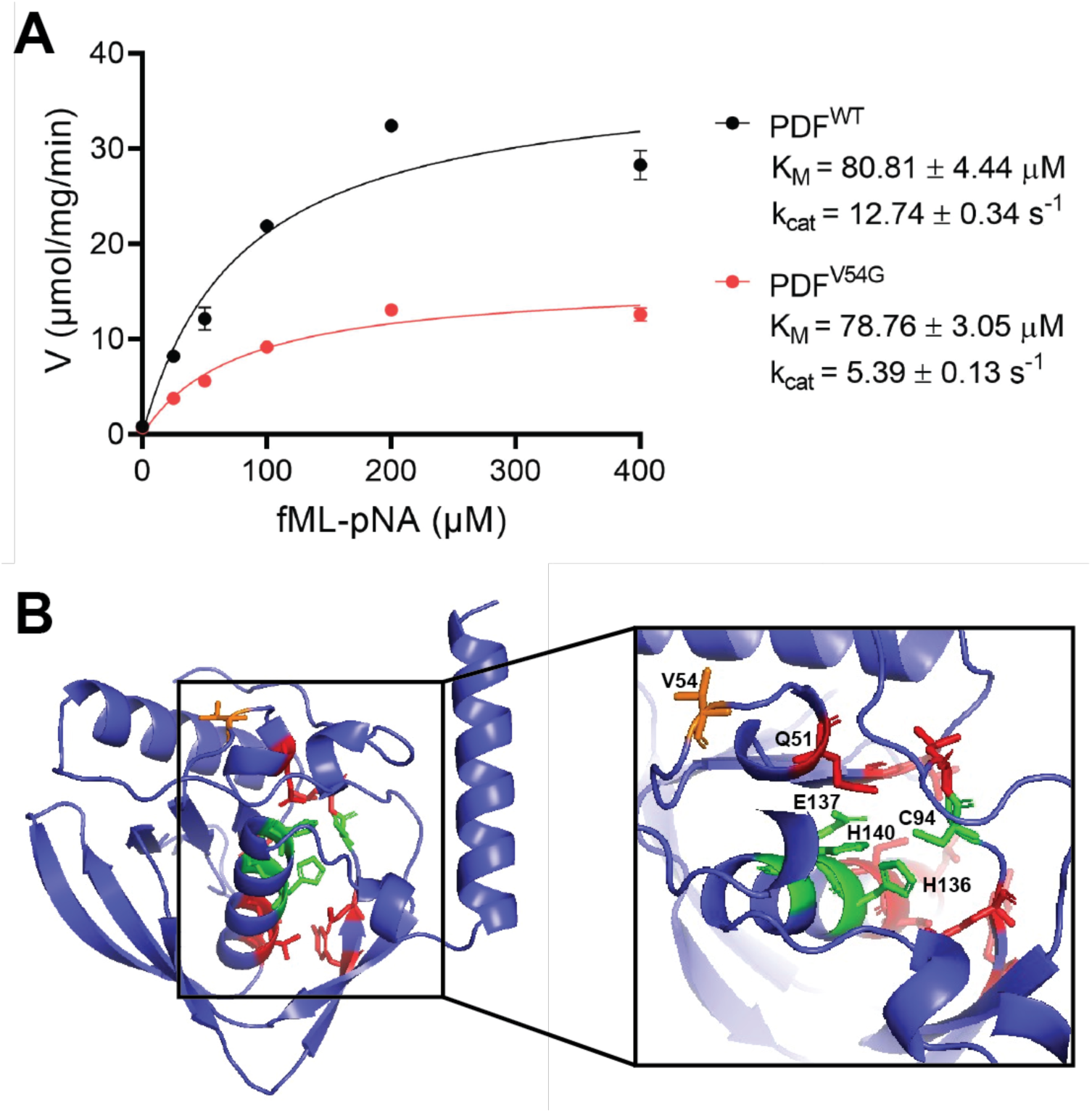
The purified PDF^V54G^ enzyme has less *in vitro* activity than PDF^WT^. A) The velocity vs substrate concentration plot for the deformylation of fML-pNA with PDF^WT^ and PDF^V54G^. Initial rates were calculated from reaction curves at ≤ 60 sec. The curve was fit to the data according to *V = Vmax · [S]/(KM + [S]).* Error bars represent the standard deviation of the mean for three replicates. B) A structural model of PDF^WT^ predicted with AlphaFold(33, 34). Conserved active site residues are green, and conserved residues known to bind the peptide substrate are red. The V54 residue is orange.

We generated a structural model of PDF^WT^ with AlphaFold (33, 34) to investigate how the V54G substitution decreases deformylase activity (Fig. 4B). There is high confidence in the accuracy of this model, with 156 residues (91.2%) having predicted local-distance difference test (pLDDT) scores greater than 90 (Fig. S2). Additionally, the model structurally aligns with the architecture of other PDF enzymes, which are highly conserved across the bacterial domain (Fig. S3). In the active site cleft, PDF enzymes have a highly conserved HEXXH metal binding motif at the base of the active site and a peptide binding pocket (35, 36). The V54 residue is in a flexible region on the surface of the enzyme and is not located in either of these conserved motifs, but similar, bulky hydrophobic amino acids are broadly conserved in this position across diverse Gram-negative bacteria (Fig. S3A). The closest fully conserved residue is Q51 (7.2 Å), which stabilizes the carbonyl oxygen of the formylmethionine substrate through hydrogen bonding (36). Given that the PDF^V54G^ variant has the same KM as PDF^WT^, it is unlikely that the V54G substitution interferes with substrate binding. Rather, the glycine substitution likely disrupts local packing interactions, increasing conformational flexibility in the loop. Given the induced-fit model that peptide deformylase uses, this increased flexibility may destabilize the catalytically competent state and reduce catalytic efficiency.

### The def^evo^ allele has pleiotropic effects on the transcriptome

To gain further understanding of the physiological impact of the *def^evo^*allele, and to potentially identify differentially expressed genes (DEGs) that would explain the increased resistance to formaldehyde, we performed RNA sequencing on exponentially growing cells. We examined transcriptomic differences in minimal medium with 15 mM methanol with and without exogenous formaldehyde treatment.

First, pairwise comparisons of log-transformed counts of WT versus the *def^evo^* mutant were performed to identify gene expression differences conferred by the *def^evo^* mutation in unstressed conditions. The *def^evo^* mutant has 820 DEGs (identified by imposing a cutoff for the false discovery rate (FDR) less than 0.01) with respect to WT, representing over 16% of the total genome (Fig. 5A, Table S1). Of these 820 DEGs, 539 genes were upregulated, and 281 were downregulated (Table S1-S3). Upregulated genes were enriched for those involved in the biosynthesis of thiamine phosphate, terpenoids, lipid A, cobalamin, and amino acids. Genes involved in tRNA charging, formaldehyde oxidation, and NADH metabolism were also upregulated in the *def^evo^*strain (Table S2). Downregulated genes fall primarily into the following categories: cell envelope physiology, phasins, Fe-homeostasis, repeats-in-toxin (RTX) exoproteins, and electron transport chain (Table S3). These results demonstrate that the *def^evo^* mutation has significant pleiotropic effects on the transcriptome, many of which are consistent with previously published reports demonstrating that PDF inhibition modulates membrane physiology, energy metabolism, and cellular redox conditions (27).

**Figure 5:**
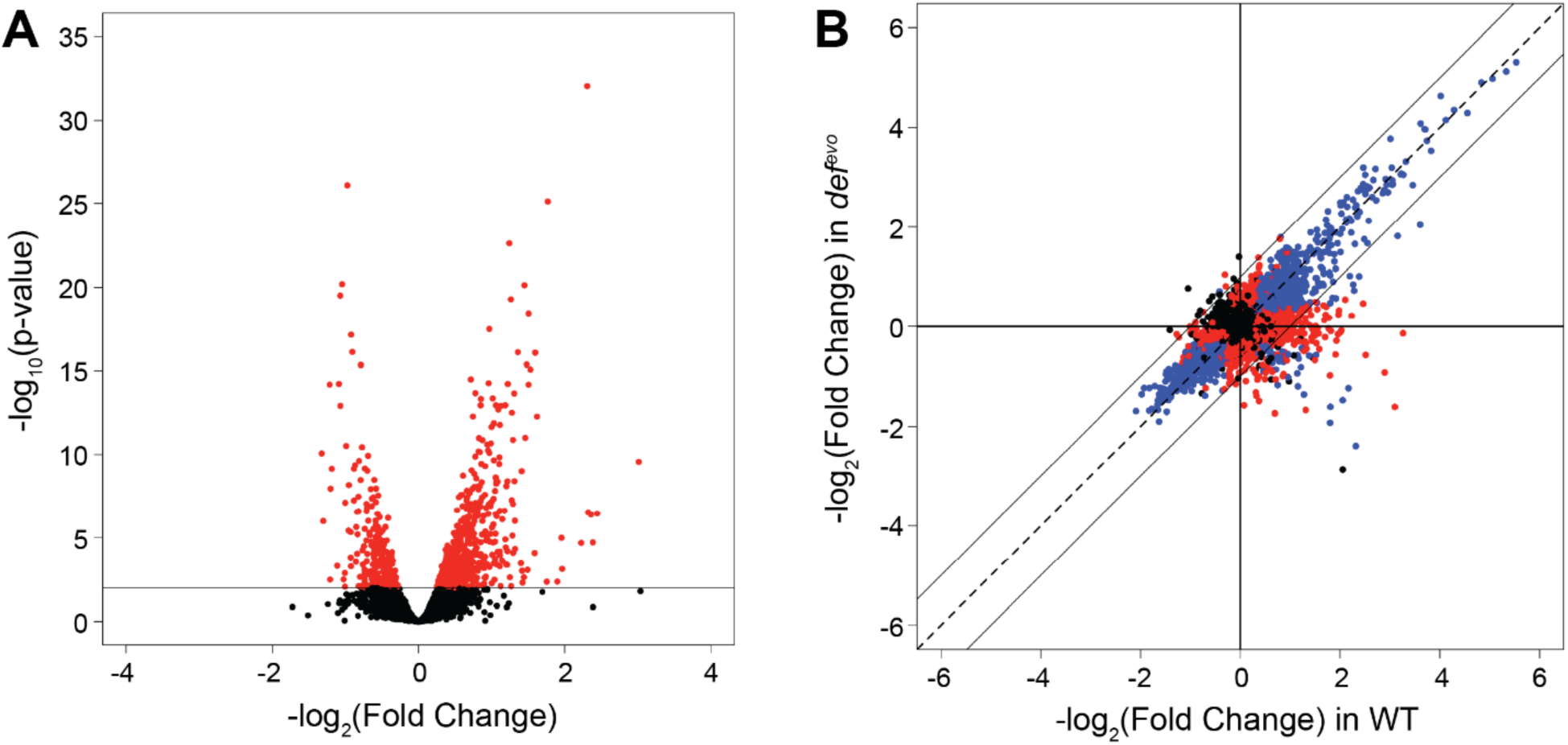
Lowering PDF activity has pleiotropic effects on the transcriptome. A) The *def^evo^* mutant differentially expresses 820 genes (p adjusted < 0.01) compared to WT, 539 of which are upregulated and 281 are downregulated. B) Comparative gene expression changes of the WT and *def^evo^* strains under exposure to 5mM formaldehyde compared to an untreated condition (log2 fold changes between the conditions). Both strains demonstrate largely corresponding responses to formaldehyde stress. Each point corresponds to a single gene. Data are color-coded according to their FDR values as follows: black points = FDR > 0.01 for both genotypes, red points = FDR ≤ 0.01 for one of the two genotypes, blue points = FDR ≤ 0.01 for both genotypes.

We also compared the response of WT and *def^evo^* strains under formaldehyde exposure (Fig. 5B, S4). Each strain was shocked by the addition of 5 mM formaldehyde and compared to the identical strain prior to exposure. Consistent with previous formaldehyde shock experiments (21), WT differentially expressed nearly 50% of the genome (2334 genes total, 1187 up, 1147 down) after 10 min of formaldehyde exposure (Fig. 5B, Table S4). The overall magnitude of the response of the *def^evo^* mutant to formaldehyde exposure was similar to the WT response (2042 genes total, 970 up, 1072 down)(Table S5). However, only a subset of these changes was common to both strains. The *def^evo^* mutant shared 60% and 73% of the upregulated and downregulated DEGs of WT, respectively, demonstrating clear differences between the two genotypes in their response to formaldehyde exposure.

Gene enrichment analysis revealed that in response to formaldehyde, both strains downregulate genes involved in the electron transport chain, central metabolism, the TCA cycle, ribosomal proteins, TonB receptors, flagellar biosynthesis, and iron metabolism. Genes involved in zeta toxins, arsenate resistance, and amino acid biosynthesis were upregulated, along with genes associated with different facets of protein biosynthesis, including tRNA processing (Tables S6, S7). Of the DEGs in WT, only 43 were oppositely regulated in *def^evo^* (Table S8), and no categories of significant enrichment were found in this subset. Gene enrichment analysis was also performed on DEGs that were unique to each strain (WT = 430 up, 311 down, *def^evo^* = 254 up, 195 down) (Table S9-S12). DEGs unique to WT’s formaldehyde response included upregulated terpenoid and porphyrin biosynthesis genes and downregulated redox-active iron metabolism genes (Tables S9, S10). Uniquely upregulated genes in the *def^evo^* strain fell into the ribonuclease Z/hydroxyacylglutathione hydrolase-like enzymatic function while downregulated genes showed enrichment for peptidoglycan synthesis, cell shape and division, and flagellar biosynthesis (Tables S11, S12).

Considering both the gene expression changes in the *def^evo^* mutant and those that are unique to its formaldehyde response, there are a number of possible explanations for the strain’s increased resistance to formaldehyde. Herein, we investigated the role of the *def^evo^*allele in protein stress and membrane physiology as it pertains to formaldehyde tolerance.

### The *def^evo^* mutant accumulates fMet-containing protein substrates

In *E. coli*, fMet has been implicated as a degron in protein quality control (25, 26). Incorrectly folded proteins are not positioned for deformylation by PDF as they exit the ribosome, and the persisting fMet recruits proteases to degrade the misfolded protein once it is released. The diminished activity of the PDF^V54G^ variant could result in an accumulation of synthesized proteins containing fMet and subsequently increase the number of proteins being targeted for degradation. We purified and treated the WT and *def^evo^* proteomes with protease to cleave all proteins into their individual amino acids and measured the relative abundance of fMet by mass spectrometry (Fig. 3C). The *def^evo^* strain indeed had 2.4-fold higher fMet levels compared to WT, demonstrating an accumulation of PDF substrates. Given the known role of fMet in protein quality control, we hypothesized that the *def^evo^*allele might mitigate formaldehyde-induced proteotoxicity by increasing the degradation of proteins damaged by formaldehyde. This hypothesis was also supported by the transcriptomics data, in which the *def^evo^* mutant upregulates genes involved in translation, including amino acid biosynthesis and tRNA charging (Table S2). Increased protein synthesis would be necessary to maintain the *def^evo^* strain’s wild-type growth rates (Fig. 2, 3, Table 1) if more proteins were being target for degradation.

### The *def^evo^* allele does not confer enhanced tolerance to thioproline

We investigated if the *def^evo^* allele impacts formaldehyde-induced proteotoxicity by comparing the growth of WT and the *def^evo^*mutant in the presence of thioproline. Thioproline can be generated from formaldehyde reactions (Fig 1B) and is a known driver of formaldehyde proteotoxicity in *E. coli* (37). If the *def^evo^* mutation confers increased formaldehyde resistance by alleviating the proteotoxic effects of formaldehyde, then the *def^evo^* strain should be more resistant to thioproline. We grew the WT and *def^evo^* strains in the presence of inhibitory levels of thioproline in three conditions with varying levels of endogenously produced formaldehyde (Fig. 6A). We assessed growth in i) succinate as a multi-carbon growth substrate (formaldehyde-free), ii) methanol as a single-carbon growth substrate (formaldehyde-generating), and iii) while cells were newly growing on methanol as they transition to methylotrophic metabolism (formaldehyde-imbalanced). We also included the τ1*efgA* mutant in these growth assays, which has elevated intracellular formaldehyde levels during the transition to methylotrophy and is predicted to experience formaldehyde-induced proteotoxicity (20). All three strains were comparably sensitive to thioproline in all conditions. Cells grown continuously in methanol were more resistant to thioproline than succinate-grown cells, but cells transitioning to methylotrophy were the most sensitive to thioproline. When undergoing a transition to methylotrophy in the presence of thioproline, the τ1*efgA* mutant no longer has its characteristic transition-induced extended lag defect (38). These results indicate that the *def^evo^* allele does not mitigate thioproline toxicity. However, further experiments revealed that thioproline may not be a formaldehyde-derived stressor in *M. extorquens* as it is in *E. coli* (Fig. S5). We therefore sought to validate if the *def^evo^* mutant mitigates formaldehyde-induced proteotoxicity through alternative methods.

**Figure 6:**
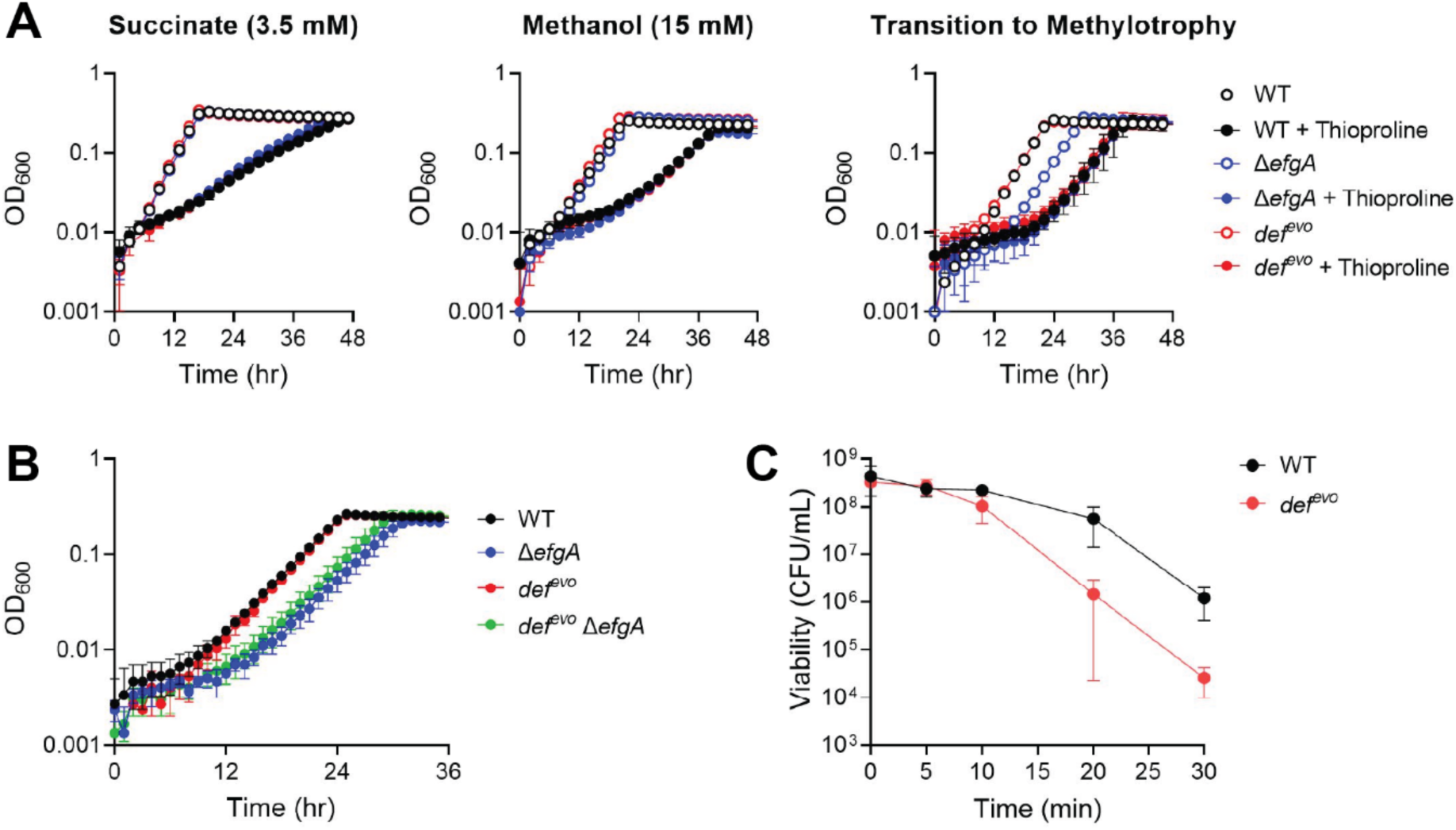
The *def^evo^* mutation does not mitigate formaldehyde-specific or general protein stress. A) WT, *def^evo^*, and Δ*efgA* strains were grown in MP with 3.5 mM succinate (+/- 15 mM thioproline), 15 mM methanol (+/- 25 mM thioproline), and 15 mM methanol during a transition to methylotrophy (+/- 5 mM thioproline). Strains have comparable sensitivities to thioproline in all conditions, although thioproline may not be a driver of formaldehyde toxicity in *M. extorquens* (Fig. S5). B) WT, *def^evo^* , Δ*efgA,* and *def^evo^* Δ*efgA* strains were grown in MP with 15 mM methanol while undergoing a transition to methylotrophy. The *def^evo^*mutation does not rescue the Δ*efgA* growth defect. C) The viability of WT and the *def^evo^* mutants after exposure to 55°C for up to 30 min. The *def^evo^* mutant has decreased viability after heat shock treatment compared to WT, indicating a potential trade-off associated with decreased PDF activity. All error bars represent the standard deviation of the mean for three biological replicates.

### The *def^evo^* allele does not mitigate proteotoxicity and has a trade off in heat shock resistance

We next evaluated growth of the Δ*efgA def^evo^* double mutant during the transition to methylotrophy. The loss of *efgA* under these conditions results in accumulation of endogenously generated formaldehyde and a loss of viability due to formaldehyde-inflicted damage (20). If the *def^evo^* mutation mitigates formaldehyde-induced proteotoxicity, then the Δ*efgA def^evo^* double mutant would have a lesser growth defect compared to the Δ*efgA* single mutant. By testing the impact of the *def^evo^* allele in these conditions, we did not constrain our query to a specific chemical species of formaldehyde-induced protein damage (e.g. thioproline). The *def^evo^* allele failed to rescue the growth defect conferred by the absence of *efgA* (Fig. 6B). These data suggested that the *def^evo^* allele may not impact formaldehyde-specific proteotoxicity and selectively alleviates exogenous, but not endogenous, formaldehyde stress.

A heat shock experiment was performed to investigate if the *def^evo^*allele confers general protein stress resistance. Exponentially growing WT and *def^evo^* strains were exposed to 55 °C for up to 30 min. We found that the *def^evo^* mutant had decreased viability after heat shock treatment compared to WT (Fig. 6C). These results demonstrate that the *def^evo^* allele does not confer a general increase in protein stress resistance and revealed an interesting tradeoff between formaldehyde resistance and resistance to heat shock. While the *def^evo^* allele confers increased resistance to formaldehyde stress, it also increases sensitivity to heat shock. We conclude that mitigating formaldehyde-specific or general proteotoxicity is not the mechanistic basis for the *def^evo^* mutant’s increased resistance to formaldehyde.

### The def^evo^ allele does not decrease formaldehyde uptake upon formaldehyde treatment

In light of the *def^evo^* mutation lacking a demonstrable benefit to protein quality control, we considered the possibility that the allele impacted formaldehyde uptake. Previous work has established that PDF activity impacts membrane integrity and permeability (27), and our transcriptomics data revealed that genes involved in membrane physiology are differentially regulated in the *def^evo^* mutant (Table S1, S12). To determine whether the *def^evo^* mutation decreased exogenous formaldehyde uptake, we employed a colorimetric assay to quantify formaldehyde depletion from the growth medium and a superfolder green fluorescent protein (sfGFP)-based formaldehyde reporter to indicate relative formaldehyde levels *in vivo*. In the plasmid encoded reporter system, sfGFP expression is under the control of the *frmR* promoter (P*frmR*) where FrmR represses *gfp* expression until a formaldehyde-derived methylene bridge deactivates FrmR. Thus, this system is effective at indicating both elevated cytoplasmic formaldehyde levels and protein damage caused by heightened formaldehyde (39).

We introduced the reporter system into WT, *def^evo^*, and Δ*efgA* strains and measured fluorescence under a range of formaldehyde concentrations (Fig. 7A). The Δ*efgA* and the *def^evo^* strains exhibited a temporal and sustained increase in cytosolic formaldehyde compared to WT. However, this difference between WT and the *def^evo^* strain does not support a model in which the *def^evo^* mutation increases formaldehyde resistance by decreasing formaldehyde uptake. Further, we do not detect any differences in rates of formaldehyde depletion from the media in the WT, *def^evo^*, and Δ*efgA* strains (WT=2.56 +/- 0.09, *def^evo^*=2.58 +/- 0.11, Δ*efgA=*2.59 +/- 0.14)(Fig. 7B).

**Figure 7:**
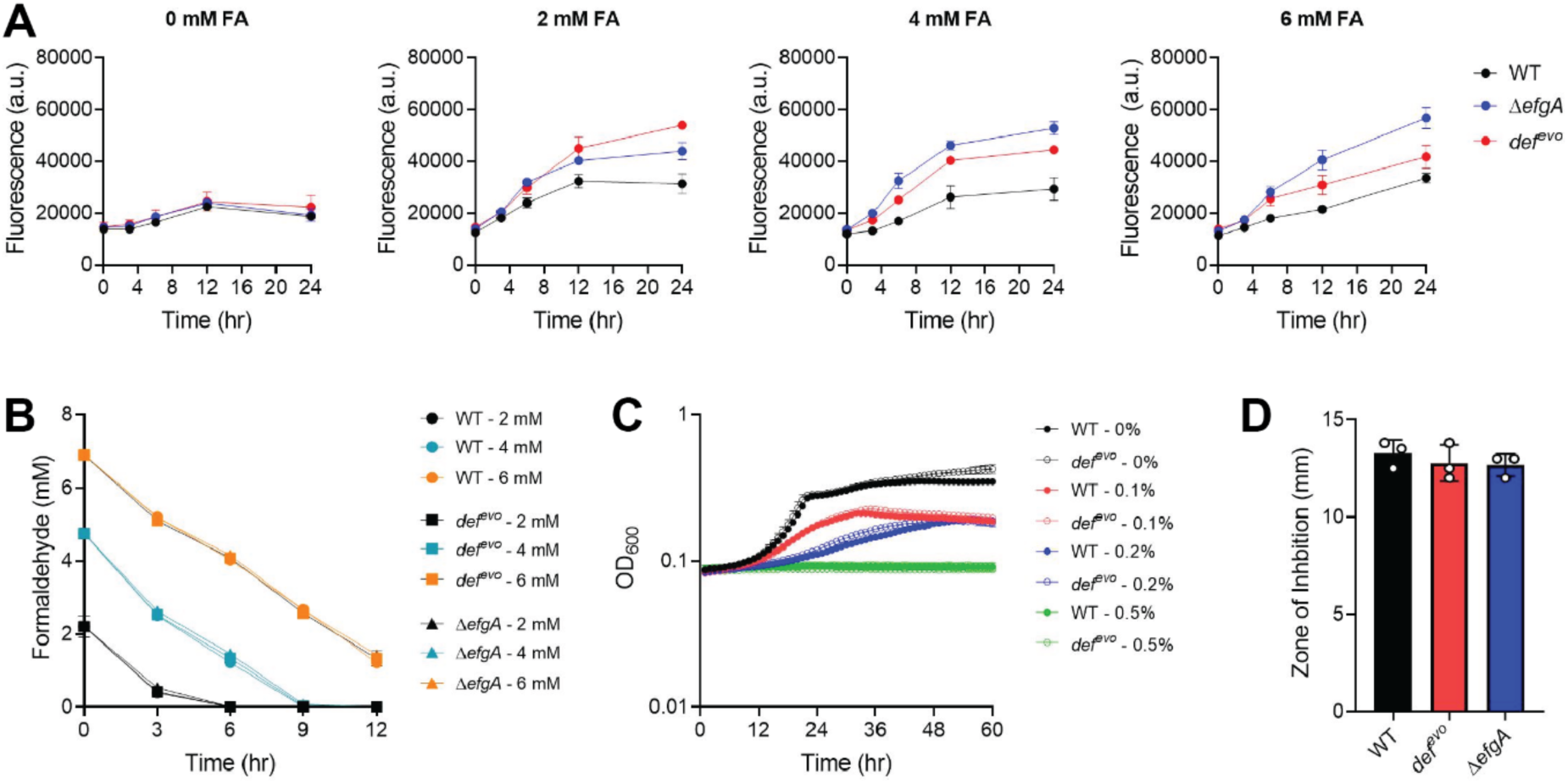
The *def^evo^* mutation does not decrease formaldehyde uptake or impact membrane permeability. A) WT, *def^evo^* , and Δ*efgA* strains were exposed to 0, 2, 4, and 6 mM formaldehyde (FA). Cytosolic formaldehyde was quantified with the P*frmR-sfgfp* reporter system. Fluorescence was measured at an emission wavelength of 510 nm with excitation at 475 nm. B) WT, *def^evo^* , and Δ*efgA* strains were exposed to 0, 2, 4, and 6 mM formaldehyde (FA). Depletion of exogenous formaldehyde was quantified with Nash assays. C) WT and *def^evo^* strains were growth in the presence of 0, 0.1, 0.2, or 0.5% butanol. D) Diffusion assays were performed by spotting 5uL of 20% SDS upon soft agar overlays of WT, *def^evo^* , and Δ*efgA* strains. Strains were equally sensitive to both membrane fluidizing agents (butanol, SDS). All error bars represent the standard deviation of the mean for three biological replicates.

Additionally, treatment with membrane fluidizing agents did not yield demonstrable phenotypic differences between WT and the *def^evo^* mutant. Both strains were equally sensitive to butanol (Fig. 7C) and SDS (Fig. 7D), suggesting that the structural integrity of the membrane is not affected by the *def^evo^* allele. We conclude that the *def^evo^* mutation does not confer increased formaldehyde resistance by decreasing formaldehyde uptake or membrane permeability.

## Discussion

Formaldehyde is a universal toxin that can cause extensive cellular damage, but is also an unavoidable product of many metabolic reactions (12–14). Cells must employ strategies to minimize biomolecular damage when formaldehyde stress is encountered, but the mechanisms governing these strategies are not well understood.

Experimentally evolving *M. extorquens* to grow on formaldehyde has allowed for the discovery of novel pathways that sense and respond to formaldehyde stress, including mechanisms for mitigating formaldehyde-induced protein damage (19). In this study, we characterized an evolved *def* allele, *def^evo^,* containing a V54G substitution. The *def* gene encodes PDF, a ribosome-associated enzyme that deformylates fMet on newly synthesized peptides. The single missense mutation in *def^evo^* allows *M. extorquens* to grow on formaldehyde as a sole carbon source, likely because the *def^evo^*strain also has increased formaldehyde resistance. We showed that the *def^evo^*strain has improved growth in the presence of formaldehyde (Fig. 2A) and that the mutant is less sensitive to formaldehyde shock in early exponential phase (Fig. 2B).

We demonstrated that the PDF^V54G^ variant has less deformylase activity and that proteins in the *def^evo^* mutant retain 2.4 times higher levels of fMet (Fig. 3, 4). We used a colorimetric assay to quantify deformylase activity of purified PDF^V54G^ and PDF^WT^ *in vitro*. Both enzymes had indistinguishable KM values of 78.76 ± 3.05 µM and 80.81 ± 4.44 µM, respectively, but the kcat of the PDF^V54G^ variant had a 58% decrease relative to PDF^WT^ (Fig. 4A). Notably, this reduction in turnover number is consistent with the quantification of fMet *in vivo.* A PDF enzyme with 58% less catalysis is expected to have ∼2.38 times higher levels of fMet substrate.

While there is strong evidence that the *def^evo^* strain has less deformylase activity than WT, it is unclear how the single V54G substitution reduces PDF activity. Comparing a structural model of the *M. extorquens* PDF to existing structures (Fig 4B, S3) revealed the V54 is not located within the highly conserved active site or the peptide binding pocket. We predict that the V54G substitution decreases turnover number by destabilizing the enzyme, but further experimentation is needed to validate this hypothesis. Additionally, although we did not observe changes in KM for the peptide substrate used in the colorimetric assay, *in vivo* PDF has thousands of different protein substrates. It is possible that the PDF^V54G^ variant has altered KM values for its natural substrates.

Transcriptomics comparing the *def^evo^* strain to WT in unstressed conditions revealed that decreasing PDF activity causes substantial changes to global gene expression (Fig. 5A). This is consistent with previous studies demonstrating that inhibition of PDF has a variety of deleterious effects on bacterial physiology, predominantly due to aberrant protein maturation and folding (27). However, the *def^evo^* mutation in *M. extorquens* positively contributes to increased formaldehyde resistance without consequence to growth rate (Fig. 2, 3, Table 1). To our knowledge, this is the first report of a reduction of PDF activity providing a benefit to any cellular physiological phenotype. There does appear to be a trade-off between formaldehyde resistance and heat resistance in the *def^evo^*mutant. We found that the *def^evo^* strain experienced more loss of viability than WT after heat shock (Fig. 6C), demonstrating the value of wild-type PDF activity under heat stress. Our results suggest that modulating PDF activity in either direction can impact cellular stress-response and may be stressor specific.

The differential expression patterns in the *def^evo^* mutant led us to pursue multiple explanations for the mutant’s increased resistance to formaldehyde. fMet is known to act as an N-terminal degradation signal (25), and we showed that more fMet persists on proteins in the *def^evo^* mutant (Fig. 3C). This finding suggests that more proteins in the *def^evo^* mutant would be targeted for degradation, and we suspected that this strain would have to increase protein synthesis to maintain its wild-type growth rate (Fig. 2, 3, Table 1). Indeed, our transcriptomics analysis revealed that the *def^evo^* mutant upregulates genes involved in translation, including amino acid biosynthesis and tRNA charging (Table S2). We postulated that this subsequent increase in protein turnover could mitigate formaldehyde-induced protein damage and cause the mutant’s increased resistance to formaldehyde. However, our investigation into the *def^evo^* allele’s effect on proteotoxicity did not definitively support a role for improved protein quality control as a mechanism of formaldehyde resistance. WT and *def^evo^*strains were not differentially sensitive to thioproline, a known driver of formaldehyde proteotoxicity in *E. coli* (Fig. 6A), and the *def^evo^* allele could not rescue the Δ*efgA* growth defect caused by endogenous formaldehyde stress (Fig. 6B). The *def^evo^* mutant also did not have a decrease in formaldehyde-induced damage via the P*frmR* reporter system (Fig. 7A), and its decrease in PDF activity did not protect the cells from general protein stress/heat shock (Fig. 6C).

Notably, work with a Δ*pepP* strain lacking the Xaa-Pro aminopeptidase demonstrated that thioproline is not a primary driver of formaldehyde toxicity in *M. extorquens* (Fig. S5). In *E. coli,* formaldehyde stress leads to the formation of thioproline, which causes proteotoxicity due to the generation thioproline-containing peptides (37). PepP-mediated cleavage of these peptides is thus crucial for mitigating both thioproline and formaldehyde stress. In *M. extorquens*, the role of PepP in mitigating thioproline stress was conserved, while its role in mitigating formaldehyde stress was not (Fig. S5). These data indicate that thioproline is not a driver of formaldehyde toxicity in *M. extorquens*. Further, as the loss of *pepP* increased formaldehyde resistance, our data suggest that PepP activity likely contributes to formaldehyde toxicity in *M. extorquens*, possibly by generating a toxic metabolite or by degrading protective peptides.

We also investigated the possibility that the *def^evo^*mutant increases formaldehyde resistance by changing membrane permeability or decreasing formaldehyde uptake. Previous studies have demonstrated that PDF inhibition influences membrane integrity (27), and we found that the *def^evo^* mutant differentially regulates genes involved in cell envelope physiology in formaldehyde-stressed and unstressed conditions (Table S1, S12). However, WT and *def^evo^* strains have comparable sensitivities to the membrane fluidizing agents SDS and butanol (Fig. 7C,D), and we determined that the *def^evo^* mutant has increased intracellular formaldehyde levels via the P*frmR* reporter system (Fig. 7A). These results indicate that the *def^evo^* mutation does not increase formaldehyde resistance by affecting the integrity of the membrane or by decreasing formaldehyde uptake.

While our work clearly establishes that decreasing PDF activity increases formaldehyde resistance in *M. extorquens*, we were unable to elucidate the mechanistic basis in this study. Transcriptomics revealed that *def^evo^* mutation has a myriad of effects on cellular physiology, and there could be a combination of factors that contribute to the *def^evo^* strain’s decrease in formaldehyde sensitivity. Future work will be needed to clarify the complex role of protein maturation in tolerating toxic levels of formaldehyde.

## Materials and Methods

### Bacterial strains, media, and chemicals

The strains of *Methylobacterium extorquens* (40) used in this study are derivatives of PA1, in which genes for cellulose synthesis have been deleted to prevent aggregation and optimize growth measurements in liquid media (41–43). Therefore, the genotype referred to as “wild-type” (CM2730) in this work is more accurately Δ*cel*. The *ΔefgA* strain (CM3745) has a single deletion at the locus *Mext_4158*. The *def^evo^* strain (CM3908) has a V54G substitution at the locus *Mext_1636*. All growth experiments with liquid media were performed at 30 °C with continuous shaking at 200 RPM in *Methylobacterium* PIPES (MP) medium with 3.5 mM succinate, unless otherwise stated. For growth on solid MP medium, Bacto Agar (15 g/L, BD Diagnostics) was added, and the succinate concentration was raised to 15 mM. Formaldehyde stock solutions (1 M) were prepared in a sealed tube by boiling 0.3 g of paraformaldehyde in 10 mL of 0.05 N NaOH for 20 min. The stock solutions were kept at room temperature and made fresh every two weeks. Unless otherwise stated, all other chemicals were purchased from Sigma-Aldrich.

### Plasmid construction

The *def* and *def^evo^* alleles were amplified from colonies of *M. extorquens* strains CM2730 and CM3908, respectively. The *frmR* gene was amplified from colonies of *E. coli* MG1655 while the the superfolder green fluorescent protein (sfGFP) under the control of the *frmR* promoter (P*frmR*) was amplified from pTR47m4-GFP (Addgene#: 102436). Primer sequences are available on request. Plasmid backbones (pCH07 and pET28a) were linearized by restriction digest and insertion fragments were inserted with an NEBuilder^®^ DNA Hifi Assembly reaction at 50 °C for 1 hr. Assemblies were transformed into NEB^®^ 5-alpha Competent *E. coli* (C2987).

### Conjugation of plasmids into M. extorquens strains

Cells were grown and exposed to *E. coli* strains containing relevant plasmids for triparental conjugative matings. Each mating contained three strains that were plated together and grown on nutrient agar plates for 24 hr: 1) an *M. extorquens* recipient strain (WT or *def^evo^*), 2) an *E. coli* donor strain (JB310 or JB317), and 3) an *E. coli* “helper” strain possessing plasmid pRK2073 that encodes functions for donor plasmid transfer. Alternatively, to introduce the pEB214 (sfGFP reporter system) into WT, *def^evo^*, and *ΔefgA* recipient strains, biparental conjugative matings were performed with derivatives of *E. coli* S17-1 that harbored the donor plasmids and also served as the helper strains (JB1310), as it chromosomally encodes functions for donor plasmid transfer. After incubation on nutrient agar, cell biomass was removed, washed, and plated on MPIPES plates supplemented with succinate, methylamine, and tetracycline and incubated at 30 °C. After sufficient time for colony formation passed, several colonies each were restreaked on homologous media for isolation.

### WT and def^evo^ formaldehyde treatment

Cultures of WT and *def^evo^* were started in biological triplicate in liquid MP medium containing succinate and, once saturated, subcultured (1/64) into acclimation cultures. When acclimation cultures were saturated, strains were subcultured (1/256) into MP medium with 3.5 mM succinate and 0-4 mM formaldehyde in Balch tubes. Alternatively, acclimation cultures were subcultured (1/256) into MP medium with 15 mM succinate, grown to an optical density (OD600) of 0.2, and then treated with 0-4 mM formaldehyde. OD600 was monitored regularly until cells reached stationary phase.

### WT and def^evo^ complementation

Cultures of WT and *def^evo^* harboring a C-terminal 6X-His tagged *def* or an empty pCH07 vector were started in biological triplicate in MP medium and, once saturated, subcultured (1/64) into acclimation cultures. When acclimation cultures were saturated, strains were subcultured (1/64) into MP medium with either 3.5 mM succinate or 3 mM formaldehyde as the carbon source. All media was supplemented with 12.5 µg/mL tetracycline.

### Actinonin minimum inhibitory concentration assay

Triplicate cultures of WT and the *def^evo^* strains were started in MP medium with 15 mM methanol as the carbon source. Once saturated, a 1/64 dilution was used to inoculate 5 mL of fresh media with 0-100 µg/mL actinonin dissolved in DMSO. OD600 was measured after 30 hr of growth.

### PDF expression and purification

*PDF^WT^ expression:* A 40 mL overnight culture of *E. coli* BL21 (DE3) with a pET28a vector encoding *def* with an N-terminal 6X-His tag (pCH05) was used to inoculate 4 L of LB supplemented with 50 µg/mL kanamycin. The culture was grown to an OD600 of 0.65 in 37 °C with continuous shaking at 200 RPM and induced with 0.1 mM isopropyl β-D-1-thiogalactopyranoside (IPTG). At the time of induction, 0.1 mM CoCl2 was also added to the culture to stabilize PDF and retain activity. The culture was grown for an additional 4 hr at 30 °C. Cells were harvested by centrifugation at 6,200 RPM for 8 min at 4 °C, and cell pellets were stored at -80 °C prior to purification.

*PDF^V54G^ expression:* An 80 mL overnight culture of *E. coli* BL21 (DE3) with a pET28a vector encoding PDF^V54G^ with an N-terminal 6X-His tag (pCH25) was used to inoculate 8 L of LB supplemented with 50 µg/mL kanamycin. The culture was grown to an OD600 of 0.45 in 37 °C with continuous shaking at 200 RPM and induced with 0.05 mM IPTG. At the time of induction, 0.1 mM CoCl2 was added to the culture to stabilize PDF and retain activity. The culture was grown for an additional 16 hr at 22 °C. Cells were harvested by centrifugation at 6,200 RPM for 8 min at 4 °C, and cell pellets were stored at -80 °C prior to purification.

*PDF purification:* Cell pellets were thawed and resuspended in buffer A (25 mM HEPES, pH 7.5, 70 mM NH4Cl, 30 mM KCl, 7 mM MgCl2, 0.2 mM CoCl2, 10% glycerol) with 10 mM imidazole and 1 mg/mL lysozyme, egg white (GoldBio, L-040-1). Cells were sonicated (MISONIX, XL 2000) using a ¼” (6 mm) tip for 3 cycles of 30 sec on and 30 sec off at 45% intensity, and cell debris was pelleted by centrifugation at 12,000 RPM for 15 min at 4 °C. After adding 1.5 mL and 0.5 mL of Ni-NTA resin slurry (GoldBio, H-350-25) to PDF^WT^ and PDF^V54G^ lysates, respectively, the soluble fractions were rotated at 6 RPM for 1 hr at 4 °C. Samples were centrifuged at 2,000 RPM for 10 min at 4 °C, and the resin was resuspended in buffer A with 10 mM imidazole and transferred to a 5 mL disposable column (Pierce, 29922). Once the resin settled, the column was washed with buffer A and 10 mM imidazole, and the protein was then eluted in buffer A with 250 mM imidazole. Final protein concentrations were measured using the BCA^TM^ Protein Assay Kit (Pierce, 23250).

Samples from the purification were treated with Laemmli Sample Buffer and heated for 10 min at 95 °C before running on a 12% SDS-PAGE gel at 200 V. PDF^WT^ was dialyzed into buffer without imidazole with 2 x 1 L buffer changes, and the variant was desalted through buffer exchange in a 5 kDa protein concentrator (Corning, 431482). Protein samples were stored at -80 °C. *Colorimetric PDF Assay*

Assays were adapted from Nguyen and Pei, 2008, Wei and Pei, 1997, and Ranjan et al., 2017. Reactions were performed in a 96-well polystyrene plate with a total volume of 250 µL. The concentration of peptide substrate, N-Formyl-methionylleucyl-p-nitroanilide (fML-pNA), was varied from 25 to 400 µM in buffer A with 1 mM TCEP and 1.6 U/mL *Aeromonas* aminopeptidase (AAP). The reaction was initiated by the addition of 50 ng of purified PDF and was monitored at 405 nm on a SpectraMax i3x Multi-Mode Plate Reader (Molecular Devices). Reactions were monitored for 2 min with absorbances measured every 10 sec. Initial rates were calculated from the first 60 sec of the progression curve. Reactions were performed at 28 °C.

### Quantification of fMet by mass spectrometry

Proteins were extracted from 10 mL of *M. extorquens* culture. Cells were harvested by centrifugation for 5 min at max speed and washed in 1 mL of MP. Cell pellets were stored at -80 °C before proceeding to the next steps. Cells were resuspended in 1 mL TRIzol reagent and lysed by bead beating. Cell suspensions were transferred to 2 mL screw cap tubes containing 0.1 mm beads and a single 3.2 mm bead. Samples were beat for 5 cycles of max RPM for 1 min and iced for 1 min. After incubation at room temperature for 5 min, samples were centrifuged for 1 min at max speed. 200 µL of chloroform was added to each supernatant, and samples were shaken vigorously by hand for 15 sec, incubated for 3 min at room temperature, and then centrifuged at 12,000 x g for 15 min at 4 °C. The upper aqueous phase was discarded. DNA was precipitated by the addition of 300 µL ethanol. Samples were mixed by inversion, incubated at room temperature for 3 min, then centrifuged at 4,000 x *g* for 10 min. Supernatants were transferred to a new tube with 1.25 mL isopropanol to precipitate proteins. Samples were incubated at room temperature for 10 min and then centrifuged at 12,000 x *g* for 15 min. Supernatants were discarded, and protein pellets were washed 3 times in 1 mL of 0.3 M guanidine HCL in 95% ethanol and once more in 100% ethanol. Pellets were allowed to air dry for 10 min. Proteins were resuspended in 200 µL of 1 M NaOH and incubated at 50°C until fully dissolved. The buffer was exchanged using PD MiniTrap G-25 columns and eluting with phosphate buffer. Protein concentrations were measured via BCA assay and samples were stored at -80 °C.

Purified proteins were digested by *S. griseus* protease in a 10:1 mass ratio. Samples were incubated at 37 °C for 16 hr, vacuum dried, and stored at -20 °C before submitting to University of Minnesota CMSP for MS analysis.

Amino acid standards were dissolved in methanol and diluted as needed; calibration curves were generated from 5-1000 pmol/µL. Samples were dried down via speed vac before being reconstituted in acetonitrile for analysis. Standards and samples were analyzed using a Sciex 6500+ QTrap Mass Spectrometer interfaced with an Eksigent M5 microLC using water with 0.1% formic acid as mobile phase A and acetonitrile with 0.1% formic acid as mobile phase B. The LC system was plumbed with an Acquity UPLC BEH HILIC column, 130 angstrom, 1.7 µm, 2.1 mm x 100mm, and run at a flow rate of 40 µL/min. Samples were run at a gradient of 100% B for 5 min, followed by a decrease to 10% B over a further 7.5 min after which the %B was increased to %100 over 30 seconds followed by a further five min at 100% B. Transitions were monitored for formylmethionine (178 → 84.1 m/z, 178 → 61.1 m/z, 178 → 56.1 m/z). Declustering potentials, entrance potentials, collision energies, and exit potentials are captured in Table S13. Raw data were processed using Skyline software and Excel.

### RNA sequencing

WT and *def^evo^* strains were inoculated into 2 mL MP (15 mM succinate) in biological triplicate and grown for 30 hr. Cells were subcultured into 5 mL fresh MP (15 mM methanol) and grown for an additional 48 hr. Cells were subcultured a second time into 50 mL fresh MP (15 mM methanol), when OD600 of the cultures reached 0.2, 10 mL of each cultures was harvested by centrifugation before formaldehyde treatment for transcriptomic analysis. The remaining culture was treated with 1 M formaldehyde to a final concentration of 5 mM for 10 min before harvesting cells from 10 mL of culture by centrifugation. Harvested cell pellets were stored at - 80 °C. RNA purification and sequencing was performed by Genewiz. A matrix of raw count data was generated using the CHURP pipeline (44). The matrix of raw counts was normalized, and formaldehyde treated samples were compared to pretreatment samples in R version 4.2.3 using DEseq2 (45). Gene enrichment analysis was performed using ShinyGO 0.77 (46) with a cutoff of FDR = 0.05 and categories were then ranked by fold enrichment.

### Heat shock

WT and *def^evo^* cultures in mid-exponential phase were incubated in a 55 °C water bath for up to 30 min. Cultures were serially diluted after heat shock and plated on solid MP medium. Viability was scored after 5 days of growth at 30 °C.

### Thioproline treatment

Cultures of WT and *def^evo^* were started in biological triplicate in liquid MP medium containing 3.5 mM succinate or 15 mM methanol and, once saturated, subcultured (1/64) into acclimation cultures. When acclimation cultures were saturated, strains were subcultured (1/64) into MP medium with the same carbon source and inhibiting thioproline levels (15 mM for succinate, 25 mM methanol) or succinate-grown acclimation cultures were used to inoculate 15mM methanol and 5mM thioproline for the transition to methylotrophy. OD600 was monitored regularly until cells reached stationary phase.

### Butanol treatment

Cultures of WT and *def^evo^* were started in biological triplicate in liquid MP medium containing 15 mM methanol and, once saturated, subcultured (1/64) into acclimation cultures. When acclimation cultures were saturated, strains were subcultured (1/64) into MP medium with methanol and 0.1, 0.2, 0.5% butanol. OD600 was monitored regularly until cells reached stationary phase.

### SDS treatment

Stationary-phase cultures (200 μL) in MP medium (succinate) were used to inoculate 4 mL of soft agar (0.5%) that was previously melted and then cooled to ∼50°C. Inoculated soft agar was agitated by shaking (∼10 sec) and then was overlaid to solid MP medium (succinate). Soft agar overlays were allowed to solidify at room temperature for 30 min. A 5 µL aliquot of 10% SDS in MP medium (no carbon) was spotted on top of agar plates. Plates were incubated at 30°C for ∼24 hr and then scored for diameter of resulting zones of inhibition.

### FrmR-based fluorescent reporter

Cultures of WT, *ΔefgA*, and *def^evo^* strains containing the plasmid pEB214 were grown in biological triplicate in 2 mL liquid MP media containing 3.5 mM succinate and 12.5 µg/mL tetracycline. Once saturated, these were subcultured (1/64 dilution) into 5 mL of homologous media also containing 50 µM cumate (4-Isopropylbenzoic acid, CAS Number 536-66-3, Sigma catalog number 268402) and growth at 30 °C with 250 rpm shaking for 48 hr. Optical densities of the cultures were measured in a Spec20 spectrophotometer, and an appropriate volume of cells was taken from each such that once resuspended they would be at an OD600 value of 0.2. These aliquots were centrifuged for 2 min at 8,000 rpm (6,200 x g). After removing the supernatant, the remaining pellets were reuspended in 1 mL of MP liquid media containing 12.5 µg/mL tetracycline, 50 µM cumate, no primary carbon source, and supplemented with formaldehyde at 0, 2, 4, 6, or 8 mM concentrations. A 200 µL volume of the cultures was transferred into opaque black-walled 96-well microtiter plates (Nunc, ThermoFisher Scientific catalog number 137101). Plates were sealed with parafilm, incubated at 30 °C, and measured in a SpectraMax i3x Multi-Mode Microplate Reader (Molecular Devices) periodically over a period of 24 hr. Fluorescence of sfGFP was detected via unlidded top measurements at 0.5 mm above the plates at low gain settings, with an excitation wavelength of α= 475 nm and an emission wavelength of α = 510 nm. Fluorescence values were normalized to non-fluorescent empty vector controls.

### Measuring formaldehyde in media

Formaldehyde concentrations in the culture media were measured as previously described (48). A 100-μl aliquot of culture was centrifuged (14,000 × g) for 2 min to isolate supernatant. A 100 μl of the supernatant or 100 μl of 1/10 dilution (diluent=MP medium, no carbon) was combined with 100 μl Nash reagent B, respectively, in 96-well polystyrene plates. Reaction plates were incubated (60°C and 10 min) and absorbance was read at 412 nm on a SpectraMax i3x Multi-Mode Plate Reader (Molecular Devices). Formaldehyde standards were prepared daily from 1 M formaldehyde stock solutions, and a standard curve was read alongside all sample measurements.

## Data Availability

All non-sequencing data can be found in a supplemental “All data” excel file where the data underlying each figure and the "Supplemental Tables" files where the supplemental tables are provided.

Raw Illumina read data and processed files of read counts per gene and normalized expression levels per gene have been deposited in the NCBI GEO database (accession no. TBD).

## Supporting Information

This article contains supporting information, including supplemental figures and all raw data.

## Acknowledgements

We would like to acknowledge Andrew J. Borchert, Lauren D. Palmer, and members of the Bazurto lab for reviewing the manuscript. We would also like to thank the Center for Metabolomics and Proteomics (CMSP) and Genewiz for technical services provided for mass spectrometry and RNA Sequencing.

## Funding and Additional Information

This work was supported by funding from National Institute of Health (NIH), National Institute of General Medical Sciences (NIGMS, https://www.nigms.nih.gov/) to JVB under award number 1R35GM146904-01. The funders had no role in study design, data collection and analysis, decision to publish, or preparation of the manuscript.

---

**Figure S1:**
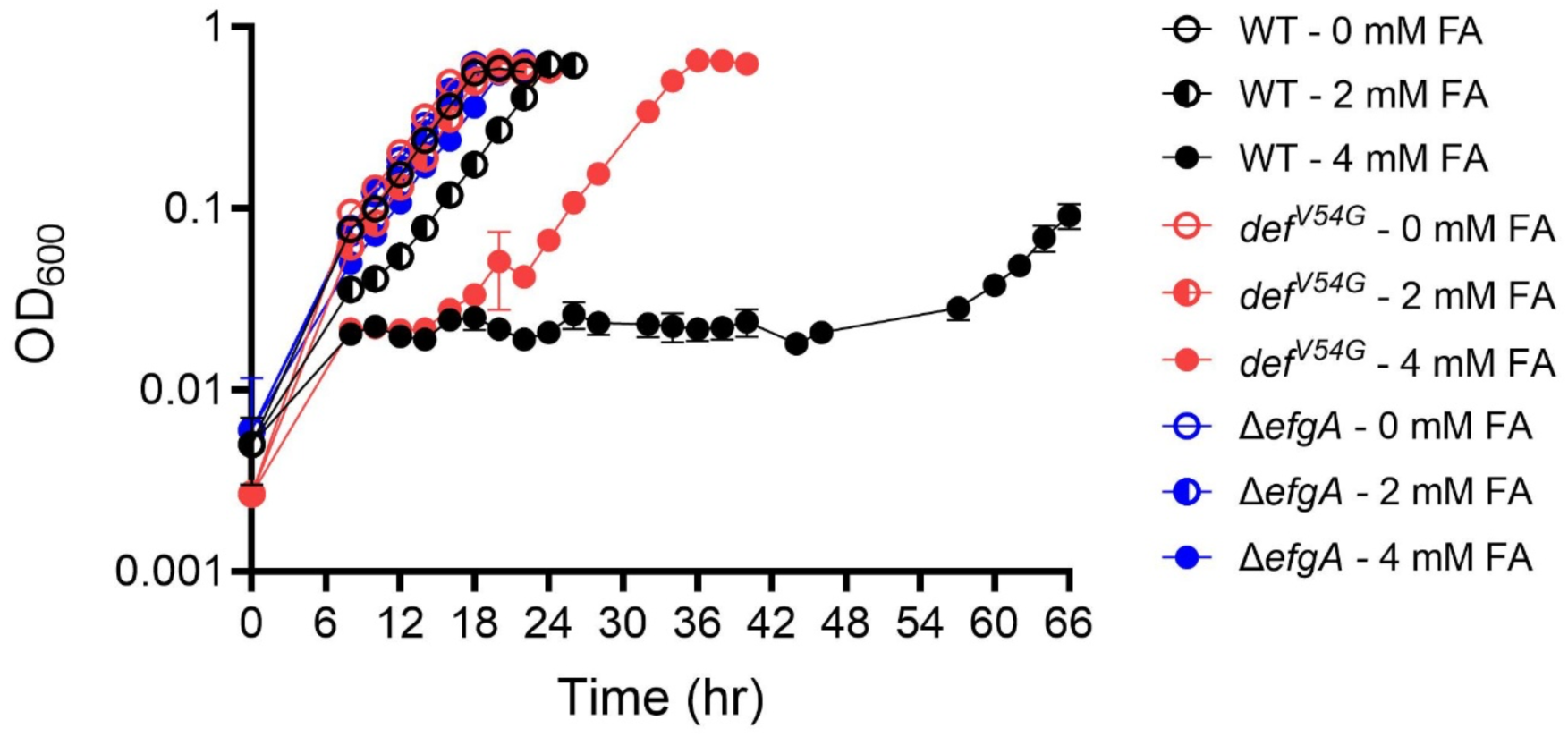
Formaldehyde growth assay with WT, *def^evo^*, and *ΔefgA*. WT, Δ*efgA*, and *def^evo^* strains were grown in MP with 3.5 mM succinate and 0, 2, or 4 mM formaldehyde. *def^evo^* and *ΔefgA* have increased resistance to formaldehyde at all concentrations tested. The error bars represent the standard deviation of the mean for three biological replicates.

**Figure S2:**
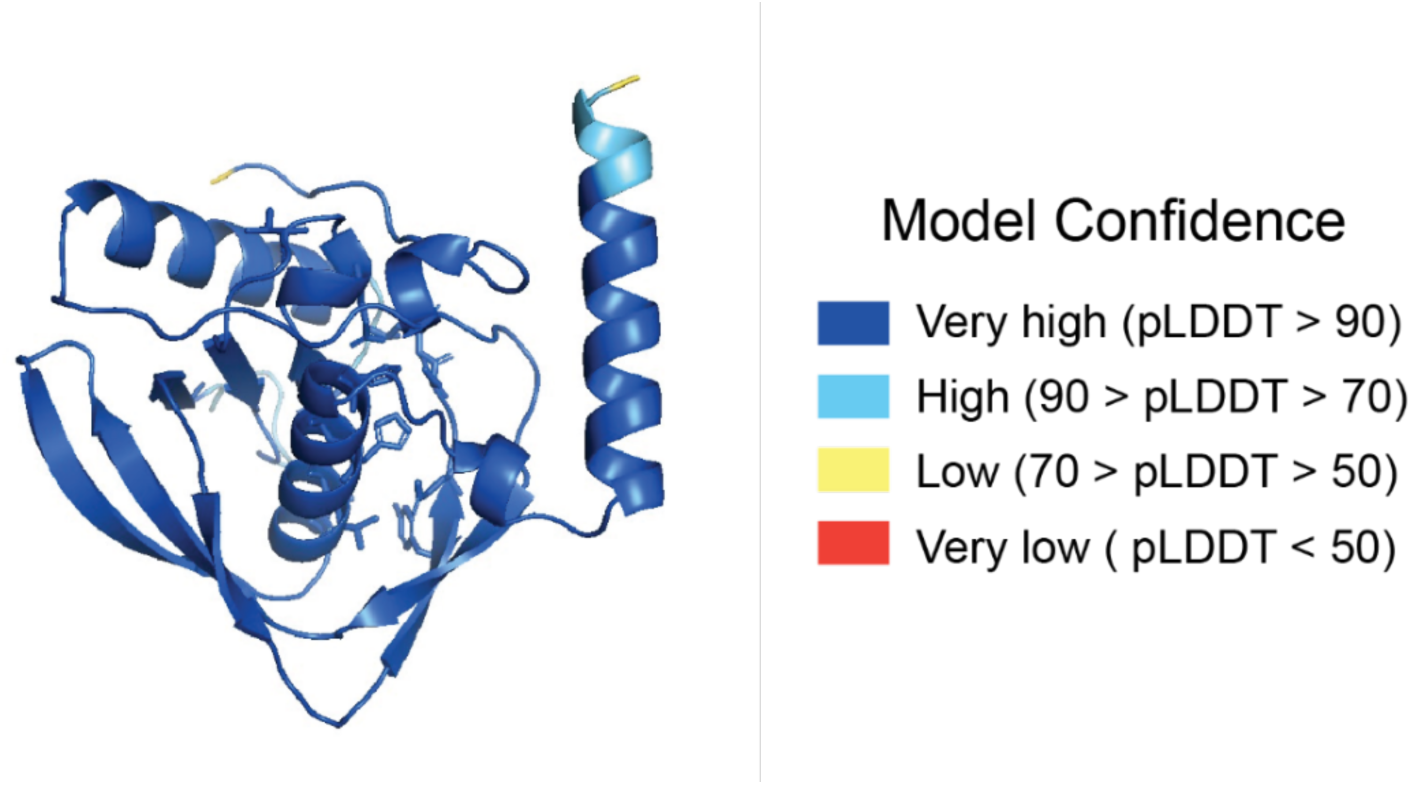
The structural model for PDF^WT^ has high confidence. The pLDDT scores reported by AlphaFold are shown. 156 of the 171 residues have scores over 90. None of the residues have scores below 50.

**Figure S3:**
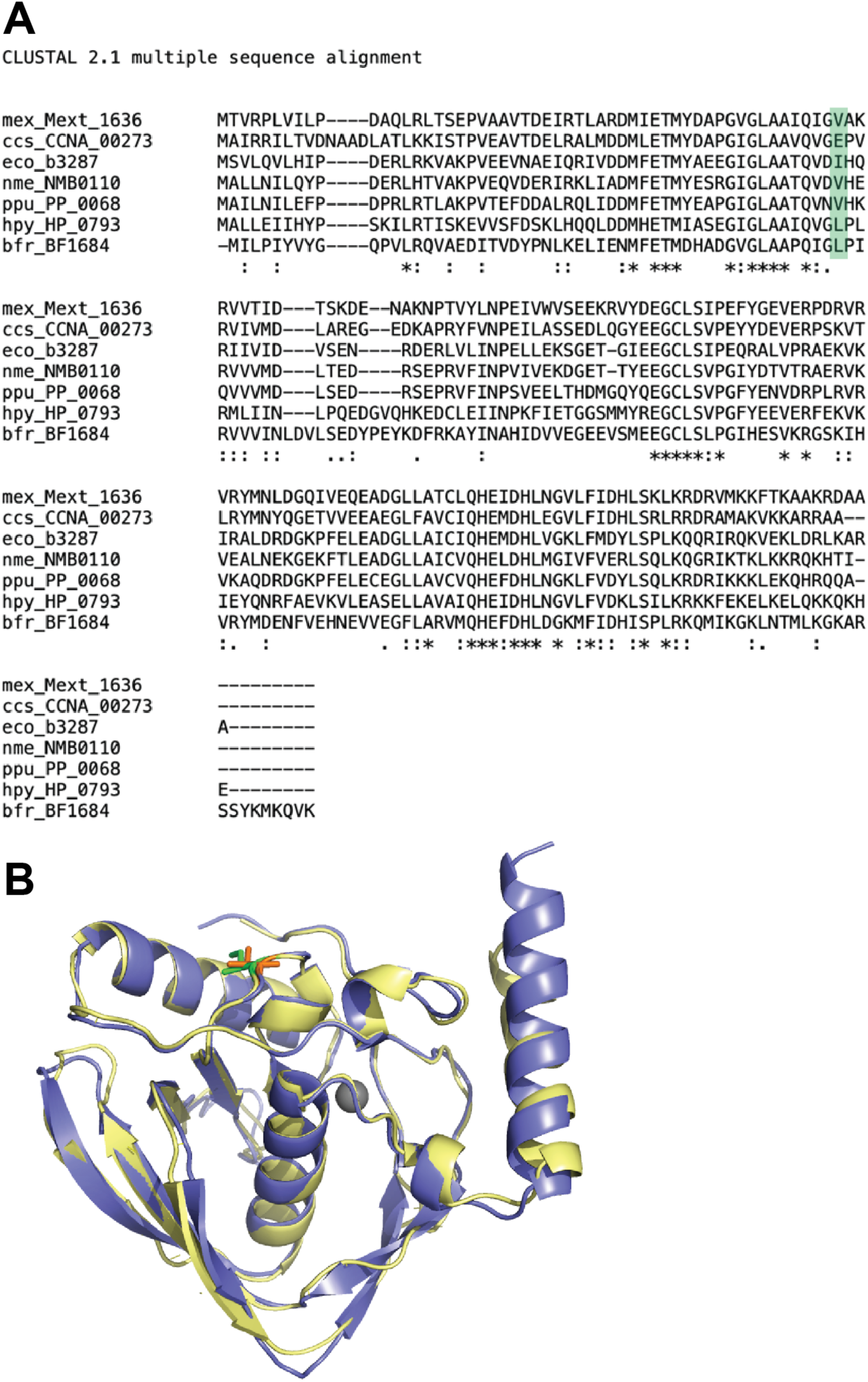
Sequence alignment of PDF from phylogenetically diverse Gram-negative bacteria. A) A Clustal Omega (47) alignment of PDF from *M. extorquens* PA1 (mex), *Caulobacter vibrioides* NA1000 (ccs), *E. coli* K-12 MG1655 (eco), *Neisseria meningitidis* MC58 (serogroup B) (nme), *Pseudomonas putida* KT2440 (ppu), *Helicobacter pylori* 26695 (hpy), and *Bacteroides fragilis* YCH46 (bfr). Conservation of residues is indicated when identical (*), strongly similar (:), or weakly similar (.). Green shading indicates position of Valine 54 in *M. extorquens.* B) *M. extorquens* PDF model (blue) aligned to solved structure of *E. coli* PDF (PDB accession: 1DFF) (yellow). The *M. extorquens* V54 residue is highlighted in orange. The *E. coli* I53 residue is highlighted in green.

**Figure S4:**
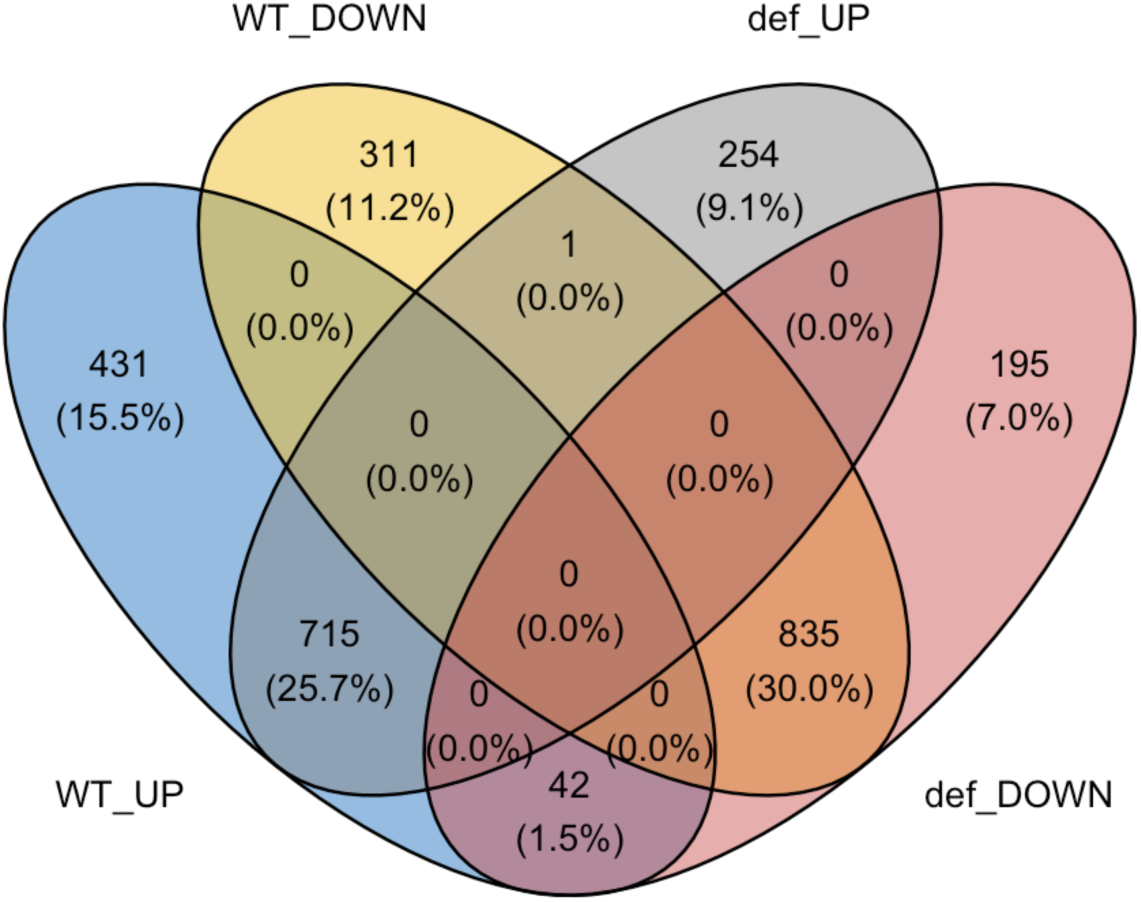
Venn diagram comparing gene expression changes of WT and *def^evo^* strains under formaldehyde shock. Comparison of differentially expressed genes in WT and *def^evo^* after exposure to 5 mM formaldehyde for 10 min. Data are for expression changes where FDR ≤ 0.01.

**Figure S5:**
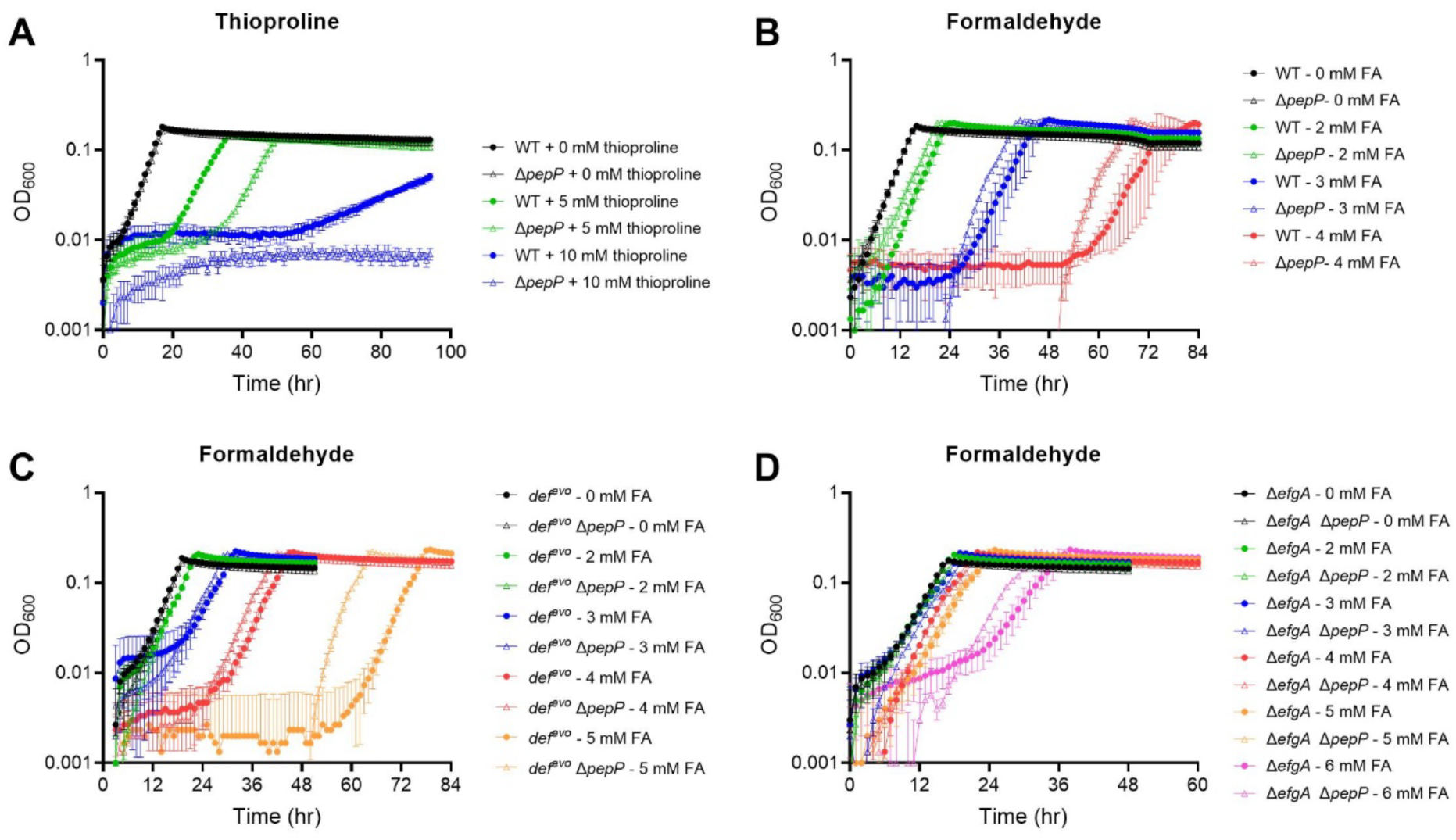
Thioproline does not drive formaldehyde toxicity in *M. extorquens*. A) WT and Δ*pepP* (JB1062, *Mext_4480*) strains were grown in MP with 3.5 mM succinate and 0, 5, or 10 mM thioproline. *pepP* encodes a Xaa-Pro aminopeptidase that hydrolyzes peptides in which the second amino acid is either proline or thioproline. In *E. coli,* PepP (EC:3.4.11.9) cleaves thioproline-containing peptides to mitigate formaldehyde-induced protein damage, and mutants lacking *pepP* have increased sensitivity to formaldehyde and thioproline (37). This data demonstrates that PepP has a conserved role for mitigating thioproline stress in *M. extrorquens*. WT and Δ*pepP* strains were grown in MP with 3.5 mM succinate and 0, 2, 3, or 4 mM formaldehyde. The Δ*pepP* mutant grew modestly better than WT in the presence of exogenous formaldehyde, suggesting that loss of PepP-mediated peptide degradation actually alleviates formaldehyde toxicity in *M. extorquens*. A similar increase in formaldehyde resistance was seen when *pepP* was deleted in C) the *def^evo^* mutant background (JB1116) and D) the Δ*efgA* mutant background (JB1117). These findings suggest that while thioproline itself is toxic in *M. extorquens*, thioproline-containing peptides are not a driver of formaldehyde toxicity.

